# RBMP2 shapes specialized membranes for CO₂ delivery in the pyrenoid condensate

**DOI:** 10.64898/2026.06.29.735335

**Authors:** Haoyu Wu, Aastha Garde, Philippe Van der Stappen, Shan He, Jessica H. Hennacy, Shixin Yang, Zhiheng Yu, Benjamin D. Engel, Martin C. Jonikas

## Abstract

Approximately one-third of global CO₂ fixation occurs in the algal pyrenoid, a condensate of the CO_2_-fixing enzyme Rubisco traversed by membranes that supply it with CO₂. In the model alga *Chlamydomonas reinhardtii*, CO_2_ is delivered by a specialized central reticulated region of these membranes. Here we define how this region is built and identify RBMP2 as the first factor required for its biogenesis. Loss of RBMP2 prevents reticulated-region formation and impairs pyrenoid-dependent growth. RBMP2 domains perform separable roles in extending membrane tubules toward the pyrenoid center and remodeling them into narrow tubules of the reticulated region. The CO_2_-releasing carbonic anhydrase CAH3 localizes not only to the reticulated region but also to helical interfaces between tubules and minitubules, which expand in some RBMP2 truncation mutants. Our findings advance the basic understanding of pyrenoid function and establish the pyrenoid as a powerful system for studying membrane remodeling within a phase-separated condensate.

## INTRODUCTION

Approximately one-third of global CO₂ fixation takes place within the pyrenoid, an organelle found in the chloroplasts of many algae and some land plants^1–8^. The pyrenoid consists of a phase-separated matrix composed primarily of Rubisco, the CO₂-fixing enzyme, and is traversed by membranes that deliver concentrated CO₂^9–14^. In the model alga *Chlamydomonas reinhardtii* (Chlamydomonas hereafter)^15^, the pyrenoid-traversing membranes (Fig. 1a) are required for growth under CO₂-limiting conditions^16–18^, underscoring their essential role in the pyrenoid CO₂-concentrating mechanism.

**Fig. 1.**
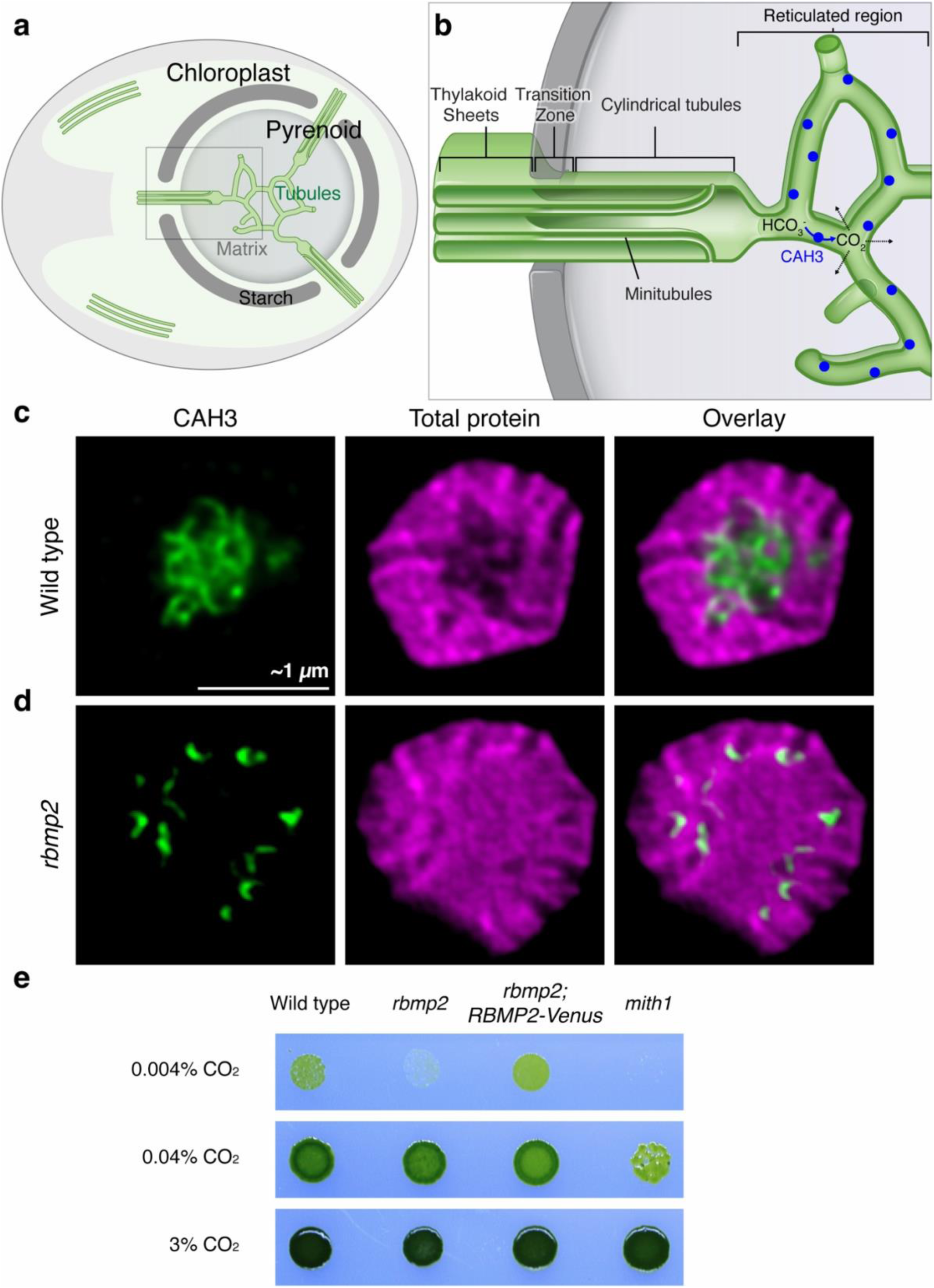
The *rbmp2* mutant mislocalizes CAH3 and has a growth defect under very low CO₂. **a,** Schematic of the Chlamydomonas pyrenoid within the cup-shaped chloroplast. The pyrenoid consists of a Rubisco-containing matrix, a surrounding starch sheath and a matrix-traversing membrane tubule network. **b,** Model of pyrenoid tubule architecture. Thylakoid sheets merge into cylindrical tubules as they enter the pyrenoid, while the inter-thylakoid stromal spaces transition into minitubules. Toward the pyrenoid center, cylindrical tubules narrow and merge into the reticulated region, where CAH3 converts bicarbonate to CO₂. Minitubules terminate by merging with cylindrical tubules before the reticulated region. **c,d,** ExM images of single pyrenoids from wild-type **(c)** and *rbmp2* **(d)** cells stained for CAH3 by immunofluorescence and for total protein by NHS-ester staining. **e,** Spot growth assay of wild type, *rbmp2*, the rescued strain *rbmp2;RBMP2-Venus* and *mith1* under the indicated CO₂ conditions. Scale bars in ExM images are corrected for the expansion factor and represent estimated true cellular dimensions.

The Chlamydomonas pyrenoid-traversing membranes are formed from thylakoid membrane sheets in the surrounding chloroplast that enter the Rubisco matrix and are reshaped into cylindrical tubules^16,18^. In this same zone, the inter-thylakoid stromal spaces are remodeled into minitubules, smaller membrane tubes within the tubules^16^ (Fig. 1b). Toward the pyrenoid center, the cylindrical tubules converge and narrow to form a central reticulated membrane network^16^. We recently found that this reticulated region hosts the key carbonic anhydrase CAH3 (encoded by *Cre09.g415700*)^19^, the enzyme thought to convert HCO₃⁻ to CO_2_ to release CO₂^20–22^, suggesting that this region functions as the central site of CO₂ release within the pyrenoid. How this membrane architecture is built remains unknown. Understanding reticulated-region biogenesis is important for engineering a pyrenoid into crops to potentially increase yields^23–25^, as correctly positioning CAH3 will require reconstructing the membrane environments that recruit or retain it. The reticulated region also provides a model for how membranes within condensates can be re-shaped to spatially organize enzyme activity^26^.

Here, we identify RBMP2 (Rubisco-Binding Membrane Protein 2, encoded by *Cre09.g416850*), as the first known factor required for reticulated-region biogenesis. We show that RBMP2 loss eliminates this central reticulated membrane network, mislocalizes CAH3 and impairs growth under very low CO₂. RBMP2’s rhodanese domain is necessary for extension of the cylindrical tubules toward the pyrenoid’s center, whereas its membrane-contact domain and transmembrane domains are required to convert cylindrical tubules into reticulated-region tubules. By characterizing RBMP2 mutants, we discover that CAH3 localizes not only to the reticulated region but also to helical interfaces between minitubules and cylindrical tubules.

Together, these findings reveal how the pyrenoid’s CO₂-delivering reticulated membrane network is built, guiding efforts to engineer pyrenoids into plants^13^ and providing an entry point for understanding how membranes are remodeled within condensates.

## RESULTS

### Loss of RBMP2 mislocalizes CAH3 and impairs growth under very low CO₂

The predicted transmembrane protein RBMP2 localizes to the reticulated region^27^, similar to the CO₂-releasing enzyme CAH3^19^, raising the possibility that RBMP2 mediates CAH3’s localization to this region. We therefore used expansion microscopy (ExM) to ask whether RBMP2 is required for CAH3 localization. Consistent with previous observations^19^, CAH3 formed a network pattern at the center of wild-type pyrenoids, supporting its localization to the reticulated region (Fig. 1c, Extended Data Fig. 1,a and Supplementary Video 1). By contrast, in the *rbmp2* mutant (LMJ.RY0402.215371, PCR verification in Supplementary Fig. 1), CAH3 no longer formed a central network and instead mislocalized as puncta and streaks near the periphery of the Rubisco matrix (Fig. 1d, Extended Data Fig. 1,b and Supplementary Video 1).

We next asked whether this CAH3 mislocalization was associated with a growth defect. Indeed, we found that the *rbmp2* mutant grew poorly under very low CO₂ (0.004%) as compared to wild type, and this defect was rescued by expression of *RBMP2-Venus* (Fig. 1e and Supplementary Fig. 2). However, the *rbmp2* growth defect at 0.004% CO₂ was less severe than that of the tubule-lacking *mith1* mutant, which may reflect more residual CO₂ release from mislocalized CAH3 in the *rbmp2* mutant than in the *mith1* mutant. We conclude that RBMP2 is required for normal CAH3 localization and for maintaining pyrenoid function under strongly CO₂-limiting conditions.

### RBMP2 is required to form the reticulated region

CAH3 could be mislocalized in the *rbmp2* mutant because it lacks a targeting factor, or because the reticulated region is defective. To characterize the integrity of the reticulated region, we first examined the total protein distribution in the pyrenoid by ExM using NHS-ester staining^19,28–30^. In wild-type cells, pyrenoid peripheral cylindrical tubules and the central reticulated region are visible as less-protein-dense regions in the more-protein-dense matrix (Fig. 2a)^19^. In contrast, the center of *rbmp2* pyrenoids was uniformly filled with protein signal at levels observed in matrix regions, suggesting that the reticulated region was absent (Fig. 2b). The less-protein-dense region in the center of the pyrenoid was restored in *rbmp2;RBMP2-Venus* cells, indicating that the defect was specifically caused by loss of RBMP2 (Fig. 2c).

**Fig. 2.**
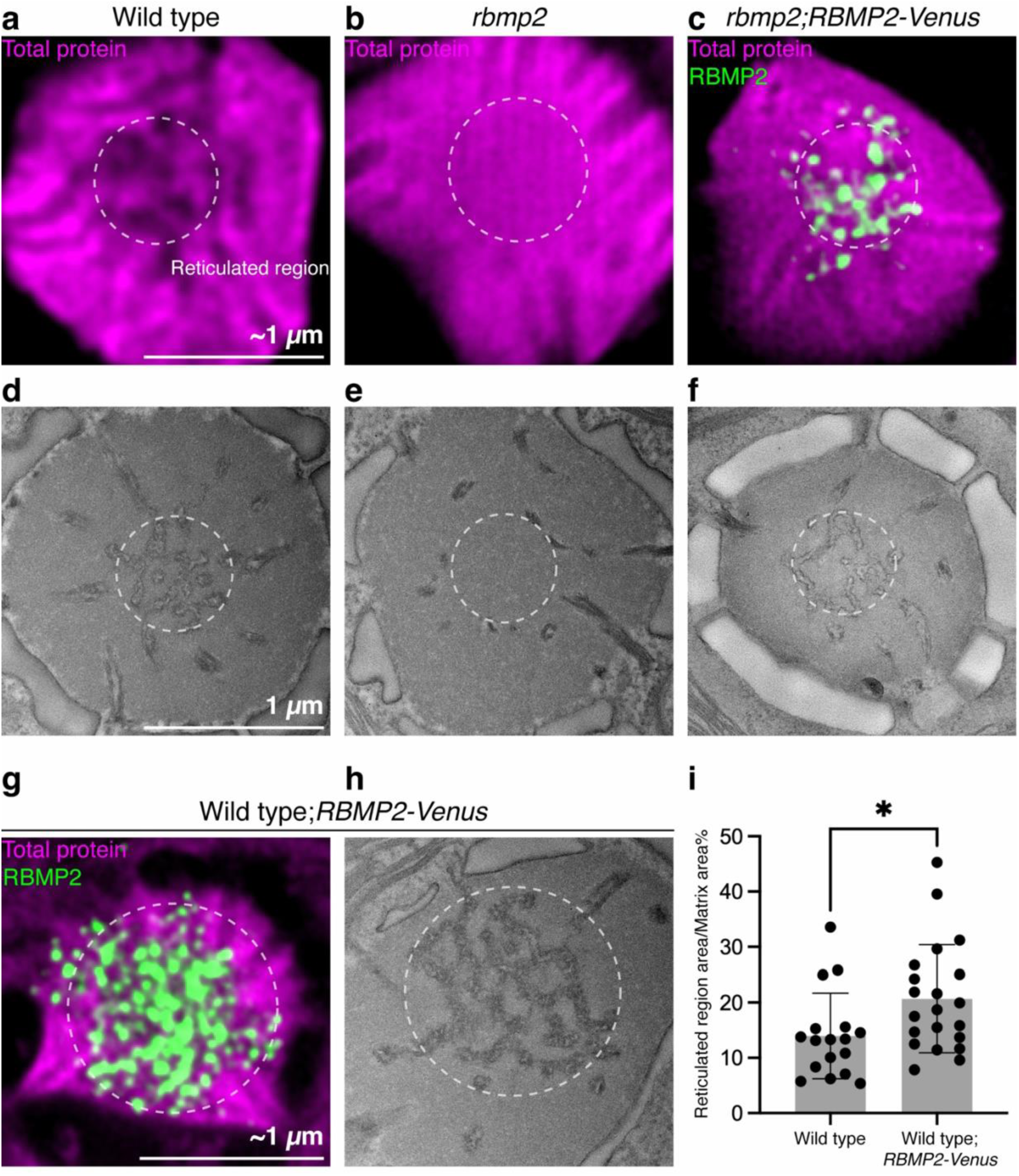
RBMP2 is required for the formation of the pyrenoid reticulated region in Chlamydomonas. **a,b,** ExM images of wild-type **(a)** and *rbmp2* **(b)** pyrenoids stained for total protein with NHS ester to visualize the pyrenoid matrix. Dashed circles indicate the center of the pyrenoid matrix where the reticulated region is visible in wild-type pyrenoids as decreased total protein staining. **c,** ExM image of the rescued *rbmp2;RBMP2-Venus* strain stained for total protein with NHS ester and for FLAG-tagged RBMP2 by immunofluorescence. **d–f,** TEM images of pyrenoids from wild type **(d)**, *rbmp2* **(e)** and the rescued *rbmp2;RBMP2-Venus* strain **(f)**. Dashed circles indicate the reticulated region or the corresponding pyrenoid center. **g,h,** ExM **(g)** and TEM **(h)** images of pyrenoids from the overexpression strain wild type;*RBMP2-Venus*. RBMP2-Venus was detected by anti-FLAG immunofluorescence in **g**. **i,** Quantification of the reticulated-region-to-matrix area ratio in wild-type and wild-type;*RBMP2-Venus* cells measured from TEM images. Scale bars in ExM images are corrected for the expansion factor and represent estimated true cellular dimensions. Error bars indicate s.d.; *=*P* < 0.05.

To definitively test whether the *rbmp2* mutant lacks a reticulated region, we examined pyrenoid ultrastructure by transmission electron microscopy (TEM). In wild-type pyrenoids, we observed cylindrical tubules that extended toward the pyrenoid center and connected to a central reticulated membrane network (Fig. 2d and Extended Data Fig. 2,a). In contrast, *rbmp2* pyrenoids contained cylindrical tubules but lacked a detectable reticulated region (Fig. 2e and Extended Data Fig. 2,d). By TEM, we observed a visible reticulated region in 7 of 52 wild-type pyrenoid cross-sections but in 0 of 52 *rbmp2* pyrenoid cross-sections (*P* = 0.0076, Fisher’s exact test). The reticulated region was restored in *rbmp2;RBMP2-Venus* cells (Fig. 2f and Extended Data Fig. 2,c); a reticulated region was observed in 5 of 50 pyrenoids. Together, these results indicate that RBMP2 is required for reticulated-region formation.

### Overexpressing RBMP2 expands the reticulated region

Having found that RBMP2 is required to form the reticulated region, we next asked whether overexpressing RBMP2 can promote the expansion of this region. In wild-type cells overexpressing *RBMP2-Venus*, we observed by ExM that RBMP2-Venus localized throughout an enlarged reticulated region (Fig. 2g). We confirmed the presence of an enlarged reticulated region in this strain by TEM (Fig. 2h and Extended Data Fig. 2,b). Quantification of TEM images showed that the reticulated-region-to-matrix area ratio increased by ∼50% in wild-type;*RBMP2-Venus* cells relative to wild type (Fig. 2i, two-sided *t*-test, *P* = 0.0232). Thus, RBMP2 is not only required for reticulated-region formation; overexpressing the RBMP2 protein is sufficient to promote the expansion of the reticulated region in wild-type cells.

### Cylindrical tubules frequently terminate prematurely in *rbmp2* mutants

Although TEM showed that the reticulated region is absent in the *rbmp2* mutant, it did not reveal the three-dimensional organization of the remaining pyrenoid membranes. We therefore used cryogenic electron tomography (cryo-ET), an approach well suited for resolving native cellular architecture and membrane organization in three dimensions^31,32^, to examine the tubule network in wild-type and *rbmp2* cells.

Consistent with previous findings^16^, by cryo-ET we observed that in wild-type cells cylindrical tubules extended from the pyrenoid periphery toward the center (Fig. 3a,b and Supplementary Video 2). As these tubules approached the pyrenoid center, their internal minitubules merged with the cylindrical-tubule membrane, thereby opening the minitubule lumen to the surrounding Rubisco matrix (Fig. 3c,d). Further in, the tubules narrowed and connected through three-way junctions to form the reticulated region (Fig. 3a–d).

**Fig. 3.**
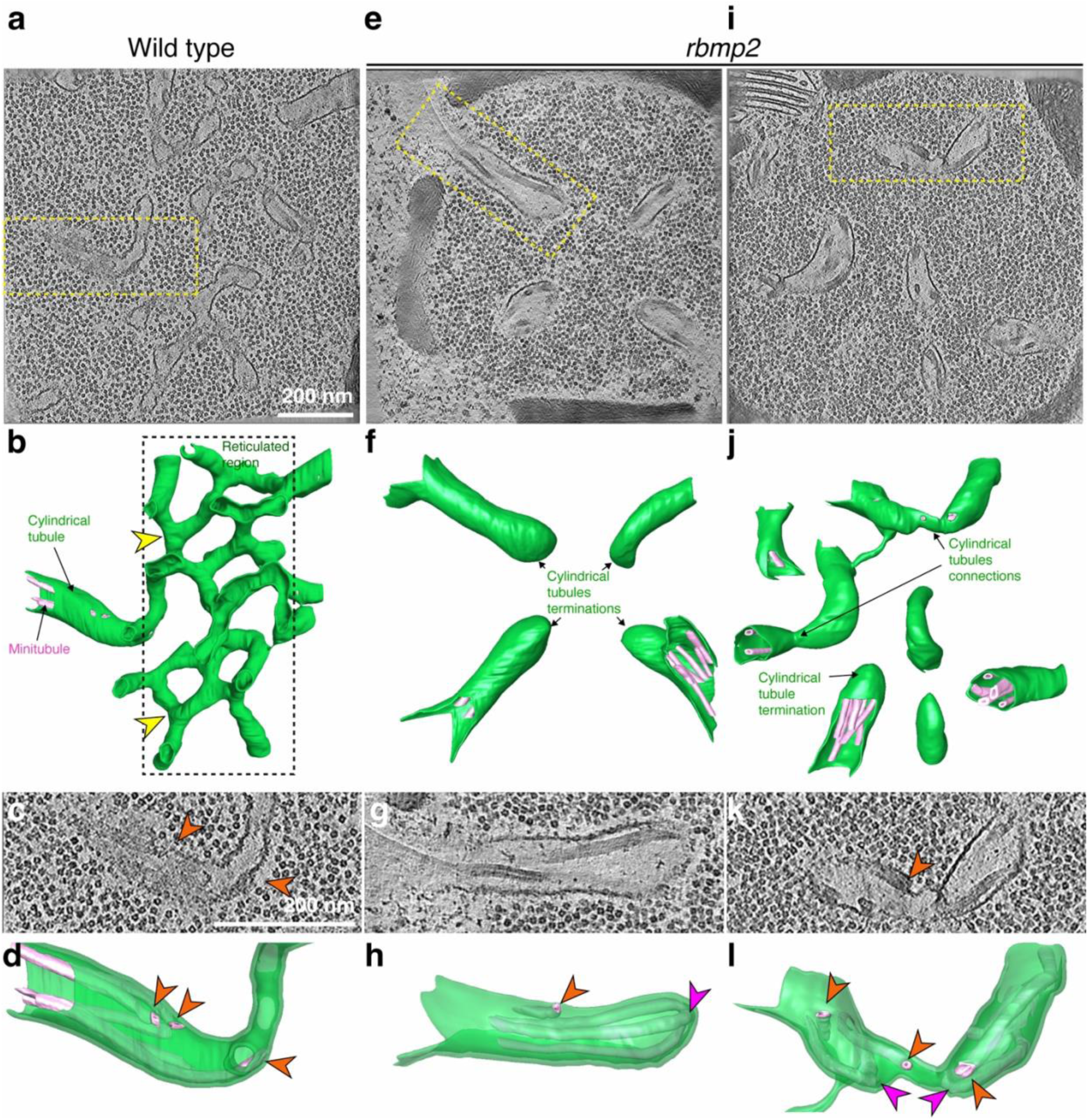
The *rbmp2* mutant forms cylindrical tubules but fails to remodel them into the reticulated region. **a,e,i,** Slices from tomograms of pyrenoids from wild-type (**a**) and *rbmp2* mutant (**e,i**) cells. Dashed boxes indicate regions shown in **c**, **g** and **k**, respectively. **b,f,j,** Segmentations of the tubule networks from the tomograms shown in **a**, **e** and **i**, respectively. Cylindrical tubules are shown in green and minitubules are shown in magenta. Yellow arrowheads indicate examples of three-way junctions between reticulated-region tubules. **c,g,k,** Magnified views of the boxed regions in **a**, **e** and **i**, respectively. Orange arrowheads indicate sites where minitubules merge with the cylindrical-tubule membrane. **d,h,l,** Magnified segmentations of the regions shown in **c**, **g** and **k**, respectively. Orange arrowheads indicate sites where minitubules merge with the cylindrical-tubule membrane. Magenta arrowheads indicate sites where minitubules merge with one another in the *rbmp2* mutant.

In the *rbmp2* mutant, cylindrical tubules still extended from the pyrenoid periphery toward the center, but we never observed them forming a reticulated network (Fig. 3e–h and Supplementary Video 2). Instead, many cylindrical tubules ended before reaching the pyrenoid center, often with closed, blunt ends (Fig. 3e,f). Consistent with this phenotype, we also observed terminated cylindrical tubules by ExM (Extended Data Fig. 1,c,d). Minitubules were still present inside terminated cylindrical tubules; some minitubules merged with the cylindrical-tubule membrane or with one another near the tubule ends (Fig. 3g,h), indicating that RBMP2 is not required for minitubule formation or fusion to the cylindrical tubules.

We also observed occasional loop-like connections between adjacent terminated tubules in the *rbmp2* mutant by ExM and cryo-ET (Extended Data Fig. 1,c,e and Fig. 3i–l). These membrane architectures suggest that some cylindrical tubules can still fuse with each other in the absence of RBMP2. Together, these results indicate that RBMP2 is required for the extension of cylindrical tubules toward the pyrenoid center and to remodel extended cylindrical tubules into the narrow, minitubule-free tubules of the reticulated region, but is not required for minitubule formation, minitubule fusion to cylindrical tubules, or tubule fusion.

### RBMP2’s rhodanese domain is required for cylindrical tubule extension

To further understand how RBMP2 builds the reticulated region, we sought to understand how its domains contribute to its functions. RBMP2 contains an N-terminal region with predicted membrane-contact probability (MCP)^33^, a central non-catalytic rhodanese domain (RHO), predicted transmembrane helices (TM) on both sides of the RHO domain and six C-terminal Rubisco-binding motifs (RBMs)^27^ (Fig. 4a and Extended Data Fig. 3,a–d). To characterize the contribution of each domain to RBMP2’s function, we transformed the *rbmp2* mutant with Venus–3×FLAG-tagged variants of RBMP2 each lacking one of these features: ΔMCP, ΔRHO, ΔTM and ΔRBM. Live-cell imaging showed that these variant proteins expressed at similar levels to the full-length protein (Extended Data Fig. 4).

**Fig. 4.**
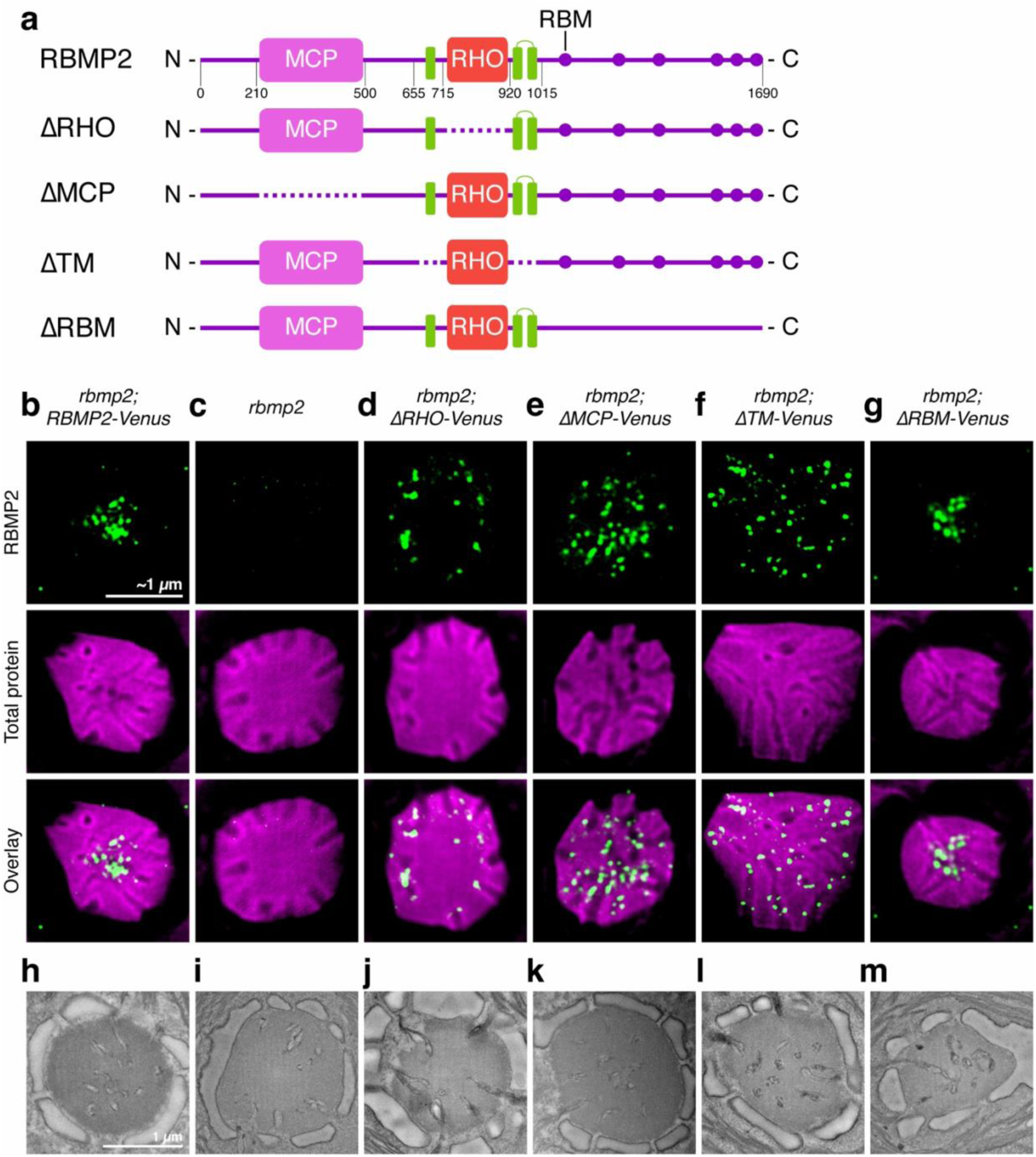
Three RBMP2 domains contribute to reticulated-region formation. **a,** Domain architecture of RBMP2 and the RBMP2 variants used in this study. RBMP2 contains an N-terminal high membrane-contact-probability region (MCP), a central rhodanese domain (RHO), a predicted transmembrane helix (TM) N-terminal to the rhodanese domain, two predicted transmembrane helices (TM) C-terminal to the rhodanese domain, and six C-terminal Rubisco-binding motifs (RBM). Dashed lines indicate the regions deleted in each RBMP2 variant. **b–g,** ExM images of pyrenoids from *rbmp2;RBMP2-Venus* **(b)**, *rbmp2* **(c)**, *rbmp2;ΔRHO-Venus* **(d)**, *rbmp2;ΔMCP-Venus* **(e)**, *rbmp2;ΔTM-Venus* **(f)** and *rbmp2;ΔRBM-Venus* **(g)** cells. Samples were stained for FLAG-tagged RBMP2 variants by immunofluorescence and for total protein with NHS ester to visualize the pyrenoid matrix. **h–m,** TEM images of pyrenoids from the strains shown in **b–g**, respectively. Scale bars in ExM images are corrected for the expansion factor and represent estimated true cellular dimensions.

We first asked whether the RHO domain is required for RBMP2 function. In *rbmp2;ΔRHO-Venus* cells, ExM showed that the reticulated region was absent and that the pyrenoid instead contained prematurely terminated cylindrical tubules, closely resembling the *rbmp2* mutant (Fig. 4c,d and Extended Data Fig. 5,c–f). TEM confirmed that *rbmp2;ΔRHO-Venus* pyrenoids lacked a normal reticulated region (Fig. 4i,j and Extended Data Fig. 2,d,e). Notably, ΔRHO-Venus still accumulated near the central ends of cylindrical tubules (Fig. 4d, Extended Data Fig. 4,b and Extended Data Fig. 5,e,f). Thus, even when the remaining portions of RBMP2 are positioned near the expected site of reticulated-region formation, loss of the RHO domain produces an *rbmp2*-like phenotype. These results suggest that the RHO domain performs an essential and upstream function in reticulated-region biogenesis, likely promoting extension of the tubules.

### MCP and TM domains are required to convert elongated cylindrical tubules into reticulated-region tubules

We next asked what role the MCP and TM domains play in RBMP2 function. In *rbmp2;ΔMCP-Venus* and *rbmp2;ΔTM-Venus* cells, ExM revealed cylindrical tubules extending into the pyrenoid center, indicating that these truncated RBMP2 proteins still supported tubule elongation (Fig. 4e,f, Extended Data Fig. 5,g–j and Extended Data Fig. 6,c). However, these extended tubules did not appear to narrow into a distinct central reticulated region, suggesting that they likely retained the morphology of cylindrical tubules. While we detected a reticulated region in 11 of 63 wild-type pyrenoids, we did not observe one in either *rbmp2;ΔMCP-Venus* cells (0 of 29 pyrenoids; *P* = 0.0115, Fisher’s exact test) or *rbmp2;ΔTM-Venus* cells (0 of 44 pyrenoids; *P* = 0.0025, Fisher’s exact test; Extended Data Fig. 6,a,b). Consistent with the potential absence of a reticulated region, ΔMCP-Venus and ΔTM-Venus localized broadly throughout the extended tubule network, rather than concentrating in a central region like full-length RBMP2 (Fig. 4e,f, Extended Data Fig. 4,c,d and Extended Data Fig. 5,g–j). TEM confirmed that the extended tubules in *rbmp2;ΔMCP-Venus* and *rbmp2;ΔTM-Venus* cells retained cylindrical-tubule morphology, rather than narrowing into reticulated-region tubules (Fig. 4k,l and Extended Data Fig. 2,f,g).

Together, these results show that RBMP2 mediates two separable functions: elongation of cylindrical tubules and their constriction into narrow, minitubule-free reticulated-region tubules. MCP and TM domains are not required for cylindrical tubule elongation. Instead, they are required to convert elongated cylindrical tubules into reticulated-region tubules.

### Deleting the six RBMs does not abolish reticulated-region formation

Finally, we asked whether the six C-terminal RBMs are required for RBMP2 function. In *rbmp2;ΔRBM-Venus* cells, we found that ΔRBM-Venus still localized to the pyrenoid center, and ExM and TEM revealed a reticulated-region-like network (Fig. 4g,m, Extended Data Fig. 2,h, Extended Data Fig. 4,e and Extended Data Fig. 5,k,l), resembling the *rbmp2;RBMP2-Venus* strain (Fig. 4b,h, Extended Data Fig. 2,c, Extended Data Fig. 4,a and Extended Data Fig. 5,a,b). By ExM, we observed the reticulated-region-like network at a frequency comparable to wild type and full-length rescue (Extended Data Fig. 6,a,b). These results suggest that the RBMs are dispensable for detectable reticulated-region formation, although they may still contribute to RBMP2 function in ways not resolved by these assays.

### MITH1 is depleted from the ends of prematurely terminated tubules in the *rbmp2* mutant

We recently found that the tubule-promoting protein MITH1^18^ forms an extended ∼100 nm Rubisco-binding coiled coil whose N-terminus binds to tubule membranes and whose C-terminus sticks away from the membranes^34^. Our data further suggested that each tubule is surrounded by a “cloud” of MITH1 coiled-coil proteins that we speculated could repel other tubules to promote proper inter-tubule spacing^34^. In the present work, those ideas led us to hypothesize that in the *rbmp2* mutant, tubule extension stops as it approaches the center of the matrix because MITH1 coats the tip of the extending tubule, repelling the tubule tip from other tubules. We further hypothesized that RBMP2 could potentially relieve this inhibition by displacing MITH1 when it binds to the tubule tips as seen in *rbmp2;ΔRHO-Venus* cells (Fig. 4d). To test these hypotheses, we investigated the localization of MITH1 in the RBMP2 truncation strains (Extended Data Fig. 7). Consistent with previous observations^19^, in wild-type cells, MITH1 localized primarily to cylindrical tubules near the pyrenoid periphery (Extended Data Fig. 7,a,b). In both *rbmp2* and *rbmp2;ΔRHO-Venus* cells, MITH1 remained associated with the peripheral portions of prematurely terminated tubules and was depleted at their tips (Extended Data Fig. 7,c–f). The depletion of MITH1 from tubule tips even in the absence of RBMP2 argues against the hypothesis that tubule extension is blocked by MITH1.

Notably, in *rbmp2;ΔMCP-Venus* and *rbmp2;ΔTM-Venus* cells, where cylindrical tubules extended toward the pyrenoid center, MITH1 remained enriched on the peripheral portions of the tubules and was less abundant along the central elongated regions (Extended Data Fig. 7,g–j). These results suggest that RBMP2’s rhodanese domain does not simply extend the tubules by expanding the MITH1-enriched region, but instead appears to function based on a different mechanism, potentially through interaction with additional factors that have yet to be discovered.

### Minitubules form helix-like patterns in the region of the cylindrical tubules closest to the pyrenoid center, which is expanded in *ΔMCP* and *ΔTM* mutants

In wild-type cells, minitubules run alongside cylindrical tubules until they approach the reticulated region, where they merge with the cylindrical-tubule membrane^16^ (Fig. 3d). Because *rbmp2;ΔMCP-Venus* and *rbmp2;ΔTM-Venus* cells lacked a reticulated region, we used cryo-ET to determine whether the morphology of minitubules was impacted in these mutants. Strikingly, minitubules were organized in an extended helix-like pattern appressed to the cylindrical tubule walls that ran along nearly their full observed length (Fig. 5e,g,i,k, Extended Data Fig. 8,i,k and Supplementary Video 4).

**Fig. 5.**
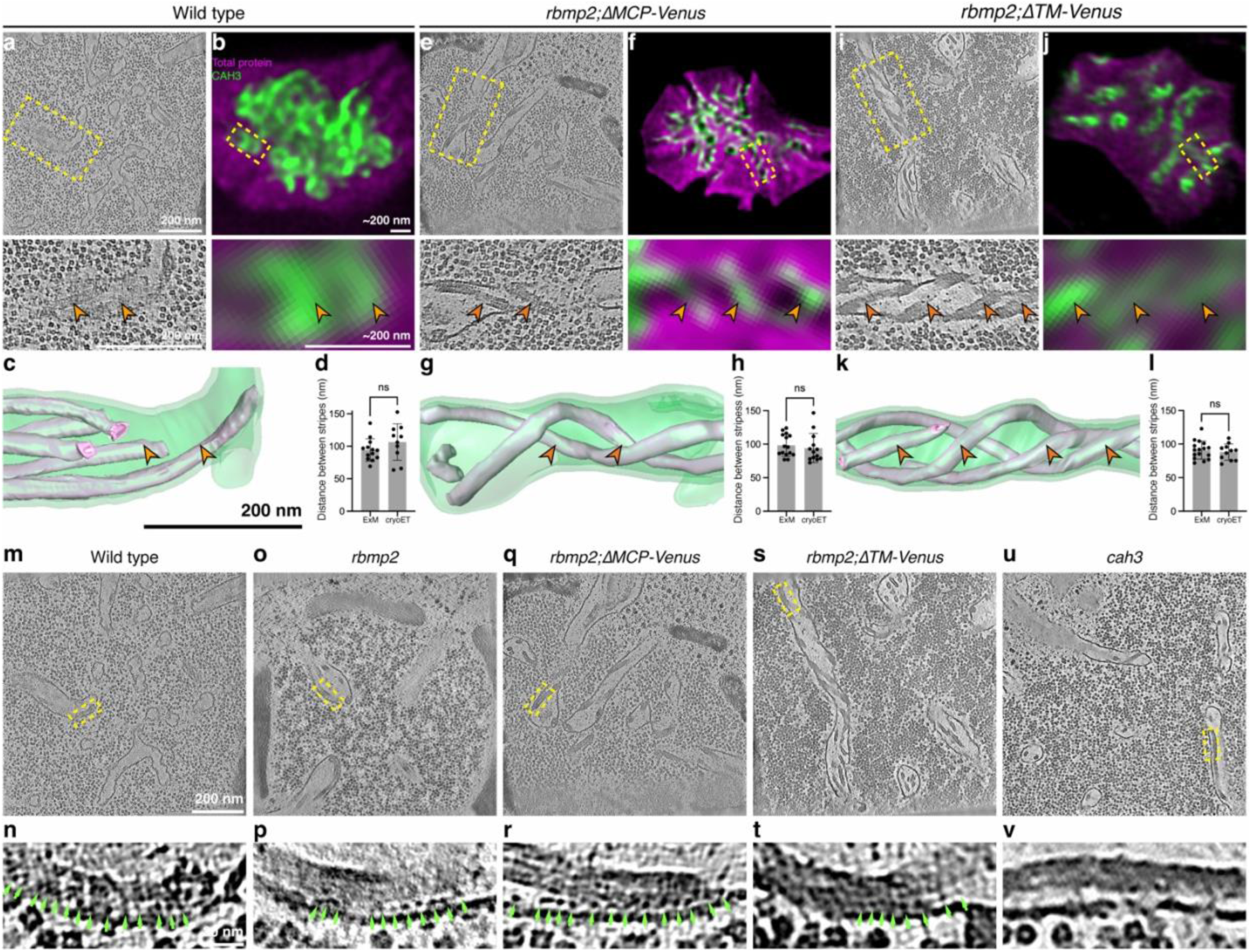
*ΔMCP* and *ΔTM* mutants have expanded helical minitubule-tubule interfaces, to which CAH3 localizes. **a,e,i,** Slices and magnified views of tomograms from wild-type (a), *rbmp2;ΔMCP-Venus* (**e**) and *rbmp2;ΔTM-Venus* (**i**). Dashed boxes indicate the magnified view regions. Orange arrowheads indicate helical minitubules. The slices shown in **a** were taken from the same tomogram as those shown in Fig. 3a,c. **b,f,j,** ExM images and magnified views of pyrenoids from wild-type (**b**), *rbmp2;ΔMCP-Venus* (**f**) and *rbmp2;ΔTM-Venus* (**j**) cells stained for CAH3 by immunofluorescence and for total protein with NHS ester. Dashed boxes indicate the magnified view regions. Orange arrowheads indicate stripe-patterned CAH3 signals. **c,g,k,** Segmentations of cylindrical tubules containing minitubules from wild-type (**c**), *rbmp2;ΔMCP-Venus* (**g**) and *rbmp2;ΔTM-Venus* (**k**) tomograms. Cylindrical tubules are shown in green, and minitubules are shown in magenta. **d,h,l,** Comparison of distances between stripes measured from CAH3 signals by ExM and between helical minitubules by cryo-ET in wild-type (**d**), *rbmp2;ΔMCP-Venus* (**h**) and *rbmp2;ΔTM-Venus* (**l**) cells. **m–v,** Slices and magnified views showing the space between minitubules and cylindrical tubules in wild-type (**m,n**), *rbmp2* (**o,p**), *rbmp2;ΔMCP-Venus* (**q,r**), *rbmp2;ΔTM-Venus* (**s,t**) and *cah3* (**u,v**) cells. Yellow dashed boxes indicate the regions magnified below. Green arrowheads indicate densities between minitubules and cylindrical tubules. The slices shown in **m,n** were taken from the same tomogram as those shown in Fig. 3a,c and Fig. 5a. The slices shown in **o,p** were taken from the same tomogram as those shown in Fig. 3e,g; those shown in **q,r** from the same tomogram as Fig. 5e; and those shown in **s,t** from the same tomogram as Fig. 5i. Scale bars in ExM images are corrected for the expansion factor and represent estimated true cellular dimensions. Data in **d,h,l** were analyzed using two-sided *t*-tests. Error bars indicate s.d.; ns, not significant (*P* > 0.05).

These results led us to look more closely at the wild-type cylindrical tubules, where we subsequently also observed such a helix-like arrangement of minitubules, but only along a short region of the tubules toward the pyrenoid center (Fig. 5a,c and Extended Data Fig. 8,a,c,d,f,g). We hereby refer to this region as the “helical-minitubule region.” By contrast, in the more peripheral portion of the cylindrical tubules, which we hereby refer to as the “non-helical-minitubule region”, the minitubules did not twist, were not all appressed to the cylindrical tubule walls, and some retained thylakoid sheet-like morphology (Extended Data Fig. 8,a,b,d,e). We conclude that in wild-type cells, cylindrical tubules have two distinct regions with different minitubule geometries, and in the *rbmp2;ΔMCP-Venus* and *rbmp2;ΔTM-Venus* mutants, the more central helical-minitubule region is expanded and extends across the center of the pyrenoid where the reticulated region would normally be.

### CAH3 localizes to the interface between the helical minitubules and cylindrical tubules

Since CAH3 normally localizes to the reticulated region, we sought to understand how the absence of this region in the *rbmp2;ΔMCP-Venus* and *rbmp2;ΔTM-Venus* mutants impacts CAH3’s localization. Unexpectedly, when we visualized CAH3 by ExM in these mutants, we observed stripe patterns along the extended cylindrical tubules (Fig. 5f,j, Extended Data Fig. 8,j,l and Supplementary Video 3). Upon closer inspection, we could also observe similar stripe patterns in some wild-type cells, at the periphery of the reticulated region (Fig. 5b and Extended Data Fig. 8,h).

The similarity of these CAH3 stripe patterns to the stripe-like appearance of minitubules in cryo-ET slices of the helical-minitubule region (Fig. 5a–l and Extended Data Fig. 8,g–l) led us to hypothesize that CAH3 localizes not only to the reticulated region but also to the minitubules in the helical-minitubule region. Consistent with this hypothesis, the spacing between helical minitubules measured by cryo-ET was similar to the spacing between CAH3 stripes measured by ExM in wild-type, *rbmp2;ΔMCP-Venus* and *rbmp2;ΔTM-Venus* cells (Fig. 5d,h,l).

To better characterize CAH3’s potential localization to the minitubules, we sought to identify CAH3 in our cryo-ET data, given the power of integrated cryo-ET and ExM for linking ultrastructure to protein identity^35^. In *epyc1* mutants, we had previously observed an unknown protein array decorating the interface between minitubules and the cylindrical-tubule membrane^36^. In the present work, we observed similar protein densities at this interface in wild-type cells and mutants (Fig. 5m–t and Extended Data Fig. 9), suggesting that these densities are not specific to the *epyc1* mutant. We thus hypothesized that these densities correspond to CAH3.

We did not observe these densities along more distal minitubules in the non-helical-minitubule region (Extended Data Fig. 9,a,b), but we observed them after minitubules moved close to the cylindrical-tubule membrane in the helical-minitubule region (Extended Data Fig. 9,c), consistent with CAH3’s localization toward the center of the pyrenoid observed by ExM (Fig. 1c, Fig. 5b and Extended Data Fig. 8,h). Furthermore, in wild-type cells, we also observed similar densities on the luminal side of the reticulated-region membrane (Extended Data Fig. 9,d, panels i–iii), where we would expect to also find CAH3 based on its localization by ExM. In the *rbmp2* mutant, which lacks the reticulated region, we observed similar cryo-ET densities near the ends of terminated cylindrical tubules, at the interface between minitubules and the cylindrical-tubule membrane (Fig. 5o,p and Extended Data Fig. 9,e,f), matching the CAH3 localization pattern observed by ExM (Fig. 1d) and further supporting the possibility that the densities were CAH3.

To more definitively determine whether these densities are CAH3, we examined the helical-minitubule region in a *cah3* mutant (LMJ.RY0402.174216) by cryo-ET. The densities were absent at the interface between minitubules and the cylindrical-tubule membrane in *cah3* cells (Fig. 5u,v and Extended Data Fig. 9,i), strongly suggesting that these densities are CAH3. Together, our observations indicate that CAH3 localizes not only to the reticulated-region membrane, but also to the interface between minitubules and cylindrical tubules in the helical-minitubule region.

## Discussion

In this study, we identified RBMP2 as the first known factor required to build the reticulated region, the membrane network at the center of the pyrenoid that houses the essential CO₂-releasing enzyme CAH3. By uncovering how this region is built, our work advances the mechanistic understanding of pyrenoid membrane morphogenesis and contributes conceptual foundations for positioning CAH3 in efforts to engineer pyrenoids into plants.

Based on our findings, we propose a mechanistic model in which reticulated-region biogenesis proceeds via two separable RBMP2-dependent steps: (1) extension of cylindrical tubules toward the pyrenoid center, and (2) conversion of cylindrical tubules into the narrow, minitubule-free tubules of the reticulated region (Fig. 6a–f and Extended Data Fig. 8,m). The phenotypes of the *rbmp2;ΔMCP-Venus* and *rbmp2;ΔTM-Venus* strains demonstrate that tubule extension can occur independently of constriction. Tubule and minitubule fusion are likely mediated by factors other than RBMP2, as indicated by the presence of loop-like cylindrical tubules and normal minitubule features in *rbmp2* cells and connected elongated cylindrical tubules in *rbmp2;ΔMCP-Venus* and *rbmp2;ΔTM-Venus* cells (Fig. 3i–l and Fig. 5e,g,i,k).

**Fig. 6.**
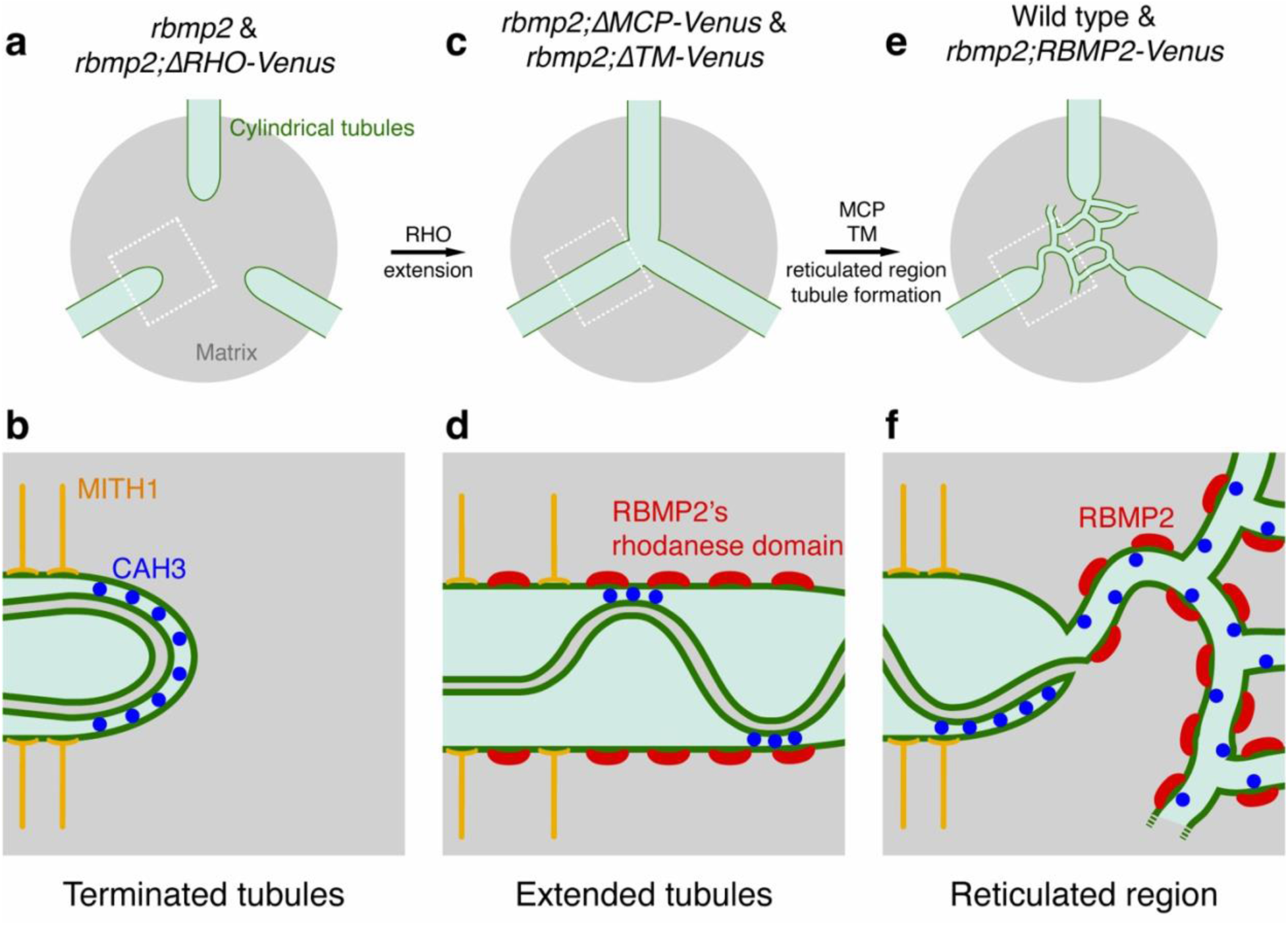
Model for how RBMP2 promotes formation of the reticulated region. **a,c,e,** Model for the roles of RBMP2 domains in reticulated-region formation. In *rbmp2* and *rbmp2;ΔRHO-Venus* cells, cylindrical tubules terminate prematurely and fail to reach the pyrenoid center **(a)**. In *rbmp2;ΔMCP-Venus* and *rbmp2;ΔTM-Venus* cells, the RBMP2 rhodanese domain promotes extension of cylindrical tubules toward the pyrenoid center, but the tubules fail to form a normal reticulated network **(c)**. In *wild-type* and *rbmp2;RBMP2-Venus* cells, full-length RBMP2 promotes both tubule extension and conversion of central cylindrical tubules into the reticulated-region tubule network **(e)**. **b,d,f,** Magnified views of the boxed regions in **a,c,e**, respectively, showing the proposed organization of MITH1, CAH3 and RBMP2 during reticulated-region biogenesis. In *rbmp2* and *rbmp2;ΔRHO-Venus* cells, MITH1 localizes along prematurely terminated cylindrical tubules, whereas CAH3 localizes near the ends of these tubules, at sites where minitubules closely approach the cylindrical-tubule membrane **(b)**. In *rbmp2;ΔMCP-Venus* and *rbmp2;ΔTM-Venus* cells, the RBMP2 rhodanese domain promotes cylindrical tubule elongation, and CAH3 localizes to expanded helical interfaces between minitubules and cylindrical tubules that span the middle of the pyrenoid **(d)**. In wild-type and *rbmp2;RBMP2-Venus* cells, full-length RBMP2 promotes further narrowing of central tubules to form the reticulated region, where CAH3 localizes to the reticulated-region membrane and to narrow helical interfaces between minitubules and cylindrical tubules **(f)**.

A notable finding of this work is that a non-catalytic rhodanese domain can influence membrane morphology, revealing a previously unrecognized role for this protein family. Non-catalytic rhodanese domains are widespread across the tree of life, but their biological functions remain poorly understood. Our results suggest that the RBMP2 RHO domain promotes cylindrical tubule elongation, pointing to a possible role in membrane-architecture control. As this domain has been proposed to mediate protein–protein interactions^37^, it may promote cylindrical tubule elongation by recruiting one or more other factors in the pyrenoid. Our previous immunoprecipitation identified LCI16 as an interactor of RBMP2^38^, and here we observed a predicted high-confidence AlphaFold co-complex structure between LCI16’s N-terminus and RBMP2’s rhodanese domain (Supplementary Fig. 3). However, the presence of a normal reticulated region in the *lci16* mutant^38^ suggests that LCI16 is not essential for reticulated-region biogenesis. Determining whether any other protein(s) act with the RBMP2 rhodanese domain to drive tubule elongation will be an important next step.

Our results suggest that the MCP and TM domains act to remodel cylindrical tubules into reticulated-region tubules. The MCP domain is predicted to be predominantly α-helical and to be in contact with the membrane, raising the possibility that it helps form or stabilize the high curvature of the reticulated region. The TM region contains predicted transmembrane helices on both sides of the RHO domain that may anchor RBMP2 to membranes, with the C-terminal helices potentially forming a membrane hairpin that may further help generate reticulated-region curvature^39^.

By characterizing RBMP2 mutants and wild-type cells, we uncovered that cylindrical tubules can be divided into two distinct regions based on the geometry of the enclosed minitubules: a “non-helical-minitubule region”, where minitubules mostly run straight, and a “helical-minitubule region”, where minitubules closely approach the cylindrical tubule wall and form helices along its surface. It will be interesting to determine in the future whether these regions defined by minitubule morphology correspond to the recently identified distinct localizations of MITH1 around the cylindrical tubules near the periphery of the pyrenoid and BST4 around the cylindrical tubules closer to the center of the pyrenoid^19^. Such a correspondence would then raise the question of what benefit, if any, the cell gains from colocalizing these proteins with the corresponding morphological features.

Localization of CAH3 near the center of the Rubisco matrix is likely crucial, as this should increase the likelihood that released CO₂ is captured by Rubisco before diffusing out of the matrix^24^. Our results suggest that CAH3 localizes not only to the reticulated region, as previously shown^19^, but additionally that it localizes to specialized portions of the cylindrical tubules, at the interface between the minitubule and the cylindrical-tubule membrane in the helical-minitubule region, which is also positioned close to the matrix center (Fig. 5m–v and Extended Data Fig. 9). This observation raises the question of what, if any, is the functional distinction between these two CAH3 localization sites, and why is CAH3 localized to both regions rather than just one? One possibility is that the two sites are specialized for different availabilities of CAH3’s substrates, H^+^ and HCO₃⁻^40,41^, which are thought to flow inward along the tubule lumen: CAH3 at the cylindrical tubule-minitubule interface likely operates with abundant substrates, whereas CAH3 in the reticulated region may operate with lower substrate concentrations, if a fraction of the substrates is depleted as they pass over the CAH3 at the cylindrical tubule-minitubule interface.

The contribution of minitubules to pyrenoid function has thus far remained elusive. Our observation that CAH3 associates with minitubules now suggests a possible role for minitubules in promoting CAH3 localization and potentially function. The extended reach of minitubules along nearly the full length of the cylindrical tubules may thus be a consequence of their contribution to CAH3 function, whose localization to the central portion of the pyrenoid appears to be necessary for efficient pyrenoid function^24^. The minitubules could potentially enhance CAH3 function by supplying CAH3 with one or both of its substrates: HCO₃⁻ and H^+^. Indeed, under some conditions, H^+^ can diffuse more rapidly along a membrane–water interface than via bulk medium^42,43^ and can be locally concentrated near a membrane^44^. It is possible that these phenomena occur in thylakoid membranes and the pyrenoid: the minitubules would thus provide additional membrane surface along which protons could diffuse to reach CAH3 in the pyrenoid, and the association of CAH3 with membranes may provide it with a higher local concentration of H^+^ (Extended Data Fig. 10,a–c).

In conclusion, our work identifies RBMP2 as a key factor mediating the biogenesis of the pyrenoid reticulated region, revealing how specialized internal membranes are constructed to position CAH3 for CO₂ release near Rubisco. The discovery that CAH3 also associates with the minitubule–cylindrical-tubule interface suggests that multiple membrane environments within the pyrenoid may contribute to efficient CO₂ delivery, and points to a previously unappreciated role for minitubules in CAH3 function. These findings contribute design principles for reconstructing pyrenoid membrane architecture in crops and establish the pyrenoid as a powerful model for membrane remodeling within phase-separated condensates.

**Supplementary Video 1**

Z-series of a wild-type (left) and an *rbmp2* mutant (right) pyrenoid combining anti-CAH3 immunofluorescence (green) with total protein stain (magenta) by ExM. Scale bars, ∼1 µm, expansion corrected.

**Supplementary Video 2**

Tomograms of a wild-type (left) and an *rbmp2* mutant pyrenoid (right) by cryo-ET. Scale bar, 200 nm.

**Supplementary Video 3**

Z-series of an *rbmp2;ΔMCP-Venus* (left) and an *rbmp2;ΔTM-Venus* (right) pyrenoid combining anti-CAH3 immunofluorescence (green) with total protein stain (magenta) by ExM. Scale bars, ∼1 µm, expansion corrected.

**Supplementary Video 4**

Tomograms of an *rbmp2;ΔMCP-Venus* (left) and an *rbmp2;ΔTM-Venus* (right) pyrenoid by cryo-ET. Scale bar, 200 nm.

## Methods

### Chlamydomonas strains and culture conditions

All strains used in this study are listed in Supplementary Table 1. Unless otherwise specified, cultures were grown to mid-log phase in Tris-acetate-phosphate (TAP) liquid medium at 22 °C in an orbital shaker (130 r.p.m.) under continuous illumination with cool white LEDs (∼200 µmol photons m⁻² s⁻¹) and ambient CO₂. Sixteen hours before each experiment, cells were pelleted at 600 × g and resuspended in Tris-phosphate (TP) medium under the same light and CO₂ conditions. Cultures were harvested for experiments at a density of approximately 2 × 10⁶ cells mL⁻¹, determined using a Countess automated cell counter.

The *rbmp2* mutant was PCR verified using three pairs of primers: an exon primer pair (CCCTTGTTCTCCTACCCCTC; GGCGTCCAAGACAAAGGTTA) that produces a 1655 bp band from wild-type template DNA, a 5′ cassette pair (CCCTTGTTCTCCTACCCCTC; GCACCAATCATGTCAAGCCT) that produces a 939-bp band from mutant template DNA, and a 3′ cassette pair (GACGTTACAGCACACCCTTG; GGCGTCCAAGACAAAGGTTA) that produces a 973-bp band from mutant template DNA.

### Spot test

Cells were cultured to mid-log phase in liquid TAP medium, pelleted at 600 × g for 5 min, and resuspended in TP medium to a final density of 6 × 10⁵ cells mL⁻¹. 10 μL of each suspension was spotted onto TP-agar plates and allowed to dry. Plates were kept in ambient air for the first 48 h before being transferred to different conditions. The plates were then incubated under 300 µmol photons m⁻² s⁻¹ illumination at different CO₂ levels (40 ppm, ambient air ∼415 ppm, or 3% CO₂). Plates were maintained under these conditions for 7 days and subsequently imaged using a Phenobooth imaging system (Singer Instruments).

### Construct design and transformation

Plasmids used in this study are listed in Supplementary Table 2. To generate Chlamydomonas strains expressing truncated variants of RBMP2, the previously described plasmid pLM155, which contains the RBMP2 genomic sequence fused to Venus-3×FLAG in the pLM005 backbone, was modified by introducing the indicated point mutations or deletions. For a higher plasmid yield, the Ori, kanR, and AphIII cassettes in pLM155 were replaced with the Ori, ampR, and AphII cassettes from pRAM118 by restriction digestion and Gibson assembly.

For transformations, Chlamydomonas cells were cultured to mid-log phase in TAP medium under ambient CO₂ using the growth conditions described above. Cells were collected by centrifugation at 600 × g for 5 min and washed twice with MAX reagent (GeneArt MAX Efficiency Transformation Reagent for Algae; Invitrogen). The final cell suspension was adjusted to 2 × 10⁸ cells mL⁻¹ in MAX reagent. For each transformation, 115 µL of cells were mixed with 5 µL of solution containing a total of ∼0.5 µg of linearized plasmid DNA (restriction enzymes are listed in Supplementary Table 2; New England Biolabs) and incubated for 5 min at 4 °C before electroporation. Electroporation was performed in a pre-cooled 2-mm gap cuvette (Bulldog Bio) using a NEPAGENE NEPA21 electroporator with the following parameters: poring pulse = 250 V, 8 ms, 50 ms interval, 2 pulses, 40% decay, + polarity; transfer pulse = 20 V, 50 ms, 50 ms interval, 5 pulses, 40% decay, ± polarity. Immediately after electroporation, 8 mL of TAP supplemented with 40 mM sucrose was added, and cultures were incubated overnight in the dark with gentle shaking. Cells were then pelleted (600 × g, 5 min) and plated on TAP-agar containing the appropriate antibiotic. After 5 days in dim light, plates were transferred to 100 µmol photons m⁻² s⁻¹ illumination for 7–14 days until colonies reached a size suitable for picking.

### Expansion microscopy

Ultrastructure expansion microscopy was performed as described previously^48^ with minor modifications. *Chlamydomonas* cells were grown in TAP medium, transferred to TP medium for 16 h, and harvested at a final density of 2 × 10⁶ cells mL⁻¹. Cells were collected by centrifugation at 600 × *g* for 5 min and resuspended in TP medium. For each sample, 200 μL of cell suspension was added to an 18-mm poly-D-lysine-coated coverslip (Neuvitro) and allowed to adhere for 10 min.

Cells were fixed for 60 min in 4% paraformaldehyde and 0.1% glutaraldehyde in 1× PBS. Fixation was quenched with 100 mM glycine for 5 min on ice, and coverslips were washed three times with cold 1× PBS. Samples were then incubated overnight at 4 °C in anchoring solution containing 2% acrylamide and 1.4% formaldehyde in bicarbonate buffer.

For gelation, monomer solution containing 10% acrylamide, 19% sodium acrylate and 0.03% *N,N′*-methylenebisacrylamide in 1× PBS was activated with TEMED and APS. The *N,N′*-methylenebisacrylamide concentration was reduced from the original U-ExM protocol to increase the expansion factor, following the TReX^49^ protocol. Activated monomer solution was added to silicone-spacer gelation chambers, and coverslips were inverted cell-side down onto the monomer solution. Gelation proceeded for 1 h on ice. Gels were then incubated in denaturation buffer containing 200 mM SDS, 200 mM NaCl and 50 mM Tris, pH 9.0, for 10 min, followed by incubation in fresh denaturation buffer at 95 °C for 90 min. Gels were washed three times with 1× PBS before staining.

For pan-protein staining, gels were incubated with 20 μg ml⁻¹ Atto 565 NHS ester in 100 mM bicarbonate buffer for 3 h at room temperature and washed three times. For immunostaining, gels were incubated overnight at room temperature in 2% BSA in PBS with anti-CAH3, 1:100 (Agrisera); anti-FLAG, 1:250 (Cell Signaling Technology); or anti-MITH1, 1:500 (Yenzyme). Gels were washed three times with 0.1% PBS-Tween 20, incubated with secondary antibodies for 3 h at room temperature, washed again with 0.1% PBS-Tween 20, and then washed three times in 1× PBS. For combined immunofluorescence and pan-protein staining, immunostaining was performed first, followed by NHS-ester staining. Gels were expanded in Milli-Q water with two water exchanges and imaged after 3 h of expansion.

### Light microscopy imaging

Light microscopy images were acquired on a VT-iSIM super-resolution spinning-disk confocal microscope equipped with a Hamamatsu Orca Quest sCMOS camera and controlled using VisiView software. Samples were imaged with a 100×, 1.35 NA silicone-immersion objective (Olympus UPLSAPO100XS) using a z-step size of 0.3 μm.

For live-cell imaging, cells were grown under the indicated conditions and transferred to an Ibidi chamber. Cells were allowed to settle for 5 min, after which excess TP medium was removed and 200 μl of 1% agarose was added to immobilize the cells. Chlorophyll autofluorescence was excited with the red laser line (594–647 nm), and Venus fluorescence was excited with the green laser line (514–532 nm).

For expanded samples, gels were first screened to identify the side containing the cells. Gels were mounted cell-side down on a 22-mm poly-D-lysine-coated coverslip in an Attofluor cell chamber. Atto 565 NHS ester was excited with the yellow laser line (561 nm). Alexa Fluor 488-conjugated secondary antibodies were excited with the blue laser line (488–491 nm), and emission was collected with a 525/25-nm filter.

### Transmission electron microscopy

Transmission electron microscopy was carried out following the previously described procedure^38^. Strains were cultured in TAP medium under ambient CO₂ to a density of approximately 5 × 10⁵ cells mL⁻¹, pelleted and resuspended in TP 16 h prior to being harvested at 1–2 × 10⁶ cells mL⁻¹. For each sample, at least 5 × 10⁷ cells were collected by centrifugation at 600 × g for 5 min in 50 mL tubes, resuspended in 1 mL TP medium, transferred to 1.5 mL screw-cap tubes, and centrifuged again under the same conditions. Pellets were fixed in 2.5% glutaraldehyde in TP medium (prepared by mixing 10 mL of 10% glutaraldehyde with 30 mL TP) and gently nutated at room temperature for 1 h. After fixation, cells were pelleted at 3,000 × g for 1 min and washed three times with 1 mL deionized water, nutating for 5 min per wash and centrifuging between washes.

Samples were post-fixed in freshly prepared osmium tetroxide solution (1% OsO₄, 1.5% w/v K₃[Fe(CN)₆], 2 mM CaCl₂) for 1–2 h at room temperature while protected from light. Because counterstaining with uranyl acetate increased background and reduced membrane contrast, only OsO₄ staining was used for all images. Following fixation, samples were dehydrated through a graded ethanol series (30%, 50%, 70%, 95%, and 100%; 1 mL each for 5 min) and two 10 min washes in 100% acetonitrile, centrifuging at 3,000 × g between steps.

Dehydrated samples were embedded in epoxy resin (34% Quetol 651, 44% nonenyl succinic anhydride, 20% methyl-5-norbornene-2,3-dicarboxylic anhydride, 2% dimethylbenzylamine catalyst; Electron Microscopy Sciences, USA) over 4 days. Embedding was initiated overnight in a 1:1 mixture of acetonitrile and uncatalyzed resin with tubes left open in a fume hood, followed by four days of nutation in catalyzed Quetol resin with daily resin replacement. On the fourth day, samples were suspended in 300–500 µL fresh resin, centrifuged at maximum speed (18,213 × g) for 20 min at 30 °C using a swinging-bucket rotor, and cured at 60–65 °C for 48 h.

Ultrathin sections (∼70 nm) were cut from cured resin blocks using a Leica Ultracut UCT ultramicrotome (Leica Microsystems, Deer Park, IL, USA), mounted on carbon film-coated 200-mesh copper grids (Electron Microscopy Sciences), and imaged at the Imaging and Analysis Center, Princeton University, using a Talos L120C G2 (S)TEM (Thermo Fisher Scientific).

### Cryo-ET sample preparation and data acquisition

Using a Vitrobot Mark 4 (FEI), cells were blotted onto carbon-coated EM grids (Quantifoil Micro Tools R 2/1, 200 Mesh, Cu; 4 μL cell culture per grid) and plunge-frozen into a liquid ethane/propane mixture at liquid nitrogen (LN_2_) temperature. The grids were blotted with Parafilm on both sides and filter paper on the back side, with blot force 6 and 8 s blot time.

Cryo-focused ion beam (cryo-FIB) milling was performed on an Aquilos 2 Cryo-Focused Ion Beam microscope (Thermo Fisher Scientific). The grids were first clipped into Autogrid supports (Thermo Fisher Scientific), and loaded onto a shuttle which was then transferred into the cryo-FIB at LN_2_ temperature^50^. The samples were first sputter-coated with Pt (30 mA, 10 Pa, 15 s), deposited an additional Pt layer using the gas injection system for 60 s (GIS, FEI), and then sputter-coated again (30 mA, 10 Pa, 15 s). Lamellas were produced using the gallium ion beam with stage tilt angles of 12°.

Tilt series of lamellae were collected on a 300-kV Titan Krios transmission electron microscope (Thermo Fisher Scientific) equipped with a cold field emission gun (cFEG), Selectris X (Thermo Scientific) energy filter, and a Falcon 4i (Thermo Scientific) camera. Using SerialEM software^51^, single-axis tilt series were recorded from −40° to +64° (in two halves separated at 12°) at 2° or 3° increments, with an object pixel size of 2.296 Å, a target defocus of −4 μm, and a cumulative electron dose of ∼150 electrons/Å².

### Cryo-ET data analysis

Image frames were aligned using MotionCor2^52^ in the RELION 5 software^53^ to correct for beam-induced motion. Tilt series were aligned and reconstructed by weighted back projection using AreTomo3^54^. Reconstructed tomograms were processed with IsoNet^55^ or DeepDeWedge^56^ for missing-wedge correction and denoising. Membrane structures in the tomograms were automatically segmented by MemBrain-seg^57^ and then manually polished in Amira. Snapshots of the tomograms were taken in IMOD^58^ and snapshots of the segmentations were taken in Dragonfly.

### Protein sequence analysis

Membrane-contact probability was predicted using the MCP predictor described previously, which estimates the likelihood that each residue contacts the lipid bilayer^33^.

Transmembrane helices in RBMP2 were predicted using DeepTMHMM with default parameters^45^. Protein-structure predictions were generated using AlphaFold2 with default parameters. For the predicted LCI16–RBMP2 complex, the N-terminal domain of LCI16 and the rhodanese domain of RBMP2 were used as input sequences, and model confidence was assessed using the predicted TM score and interface predicted TM score^59,60^. Rhodanese-domain sequences were aligned using PROMALS3D^61^. The alignment was used to compare the catalytic cysteine residue found in canonical rhodanese domains with the corresponding residue in RBMP2.

### Data availability

All data supporting the findings of this study are available within the paper and its Supplementary Information. Raw and processed microscopy data are available from the corresponding author upon reasonable request. The *Chlamydomonas* strains and plasmids generated in this study are listed in Supplementary Tables 1 and 2 and are available from the corresponding author upon reasonable request. The *rbmp2* mutant strain LMJ.RY0402.215371 was obtained from the CLiP library^46^. Protein sequence and structural analyses were performed using publicly available tools and databases as described in the Methods.

## ACKNOWLEDGEMENTS

We thank the Princeton Imaging and Analysis Center, especially John J. Schreiber and Paul Shao, for assistance with FIB-SEM and TEM experiments. We are grateful to the Cryo-EM Facility at the Janelia Research Campus, particularly Rui Yan, Momoko Shiozaki, and Xiaowei Zhao, for training and help with cryo-ET instrumentation. We thank Manon Demulder and Ricardo Righetto for training with cryo-ET instrumentation and software. We also thank the Princeton Confocal Imaging Core Facility, especially Gary Laevsky and Sha Wang, for assistance with confocal imaging. We thank Sabrina L. Ergun for providing help with biochemical assays, Jian Huang for providing guidance on protein analysis, Eric Franklin for training with cryo-FIB milling, and Victoria Crans for providing the *rbmp2* mutant strain. We further thank Ned S. Wingreen, Alistair J. McCormick, Luke Mackinder, Lianyong Wang, Qiwei Yu, Micah Burton, and members of the Jonikas lab for helpful discussions and feedback. This project was supported by the National Institute of General Medical Sciences of the National Institutes of Health under grant T32GM007388; support from the US National Science Foundation (MCB-1935444), the US National Institutes of Health (1R01GM140032-01), the US Department of Energy (DE-SC0020195) and the Howard Hughes Medical Institute to M.C.J.; and the Bill & Melinda Gates Foundation and United Kingdom Foreign, Commonwealth and Development Office (INV-054558), and the Howard Hughes Medical Institute to M.C.J.; and the Swiss Nanoscience Institute PhD School grant (P2204) and ERC consolidator grant “cryOcean” (fulfilled by the Swiss State Secretariat for Education, Research, and Innovation, M822.00045) to B.D.E. H.W. is a recipient of a fellowship from the Chinese Research Council and P.V.d.S. is funded by the Swiss Nanoscience Institute PhD School grant (P2204). The content is solely the responsibility of the authors and does not necessarily represent the official views of the funders.

## AUTHOR CONTRIBUTIONS

H.W. and M.C.J. designed the study. H.W. carried out the experiments. A.G. developed optimized ExM for the pyrenoid, and H.W. performed ExM sample preparation and imaging. A.G. first observed stripe-like CAH3 patterns in wild-type cells. H.W. first observed helix-like minitubules, stripe-like CAH3 patterns in truncation strains, and made the comparison. H.W. and S.Y. performed cryo-ET data collection. H.W. and P.V.d.S. conducted cryo-ET data analysis. S.H. and J.H.H. contributed preliminary investigations. B.D.E. and Z.Y. provided guidance in cryo-ET data collection and data analysis. H.W. and M.C.J. wrote the manuscript with input from all authors.

---

**Extended Data Fig. 1.**
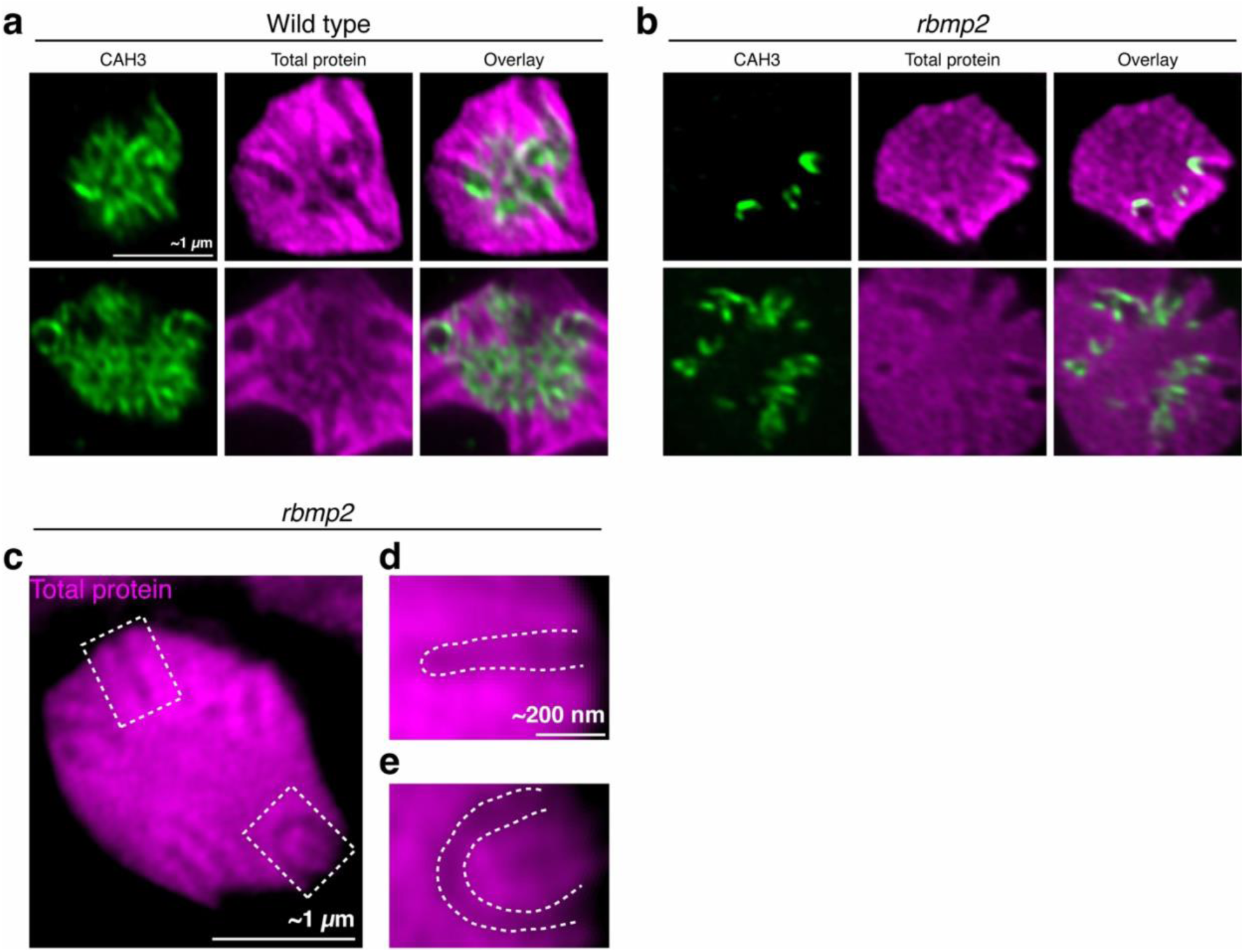
Additional ExM images of wild type and the rbmp2 mutant. **a,b,** Additional ExM images of wild-type **(a)** and *rbmp2* mutant **(b)** pyrenoids stained for CAH3 by immunofluorescence and for total protein with NHS ester to visualize the pyrenoid matrix. **c,** ExM image of an *rbmp2* pyrenoid stained for total protein with NHS ester. Dashed boxes indicate the regions shown in **d** and **e**. **d,e,** Magnified views of the boxed regions in **c**, showing a terminated cylindrical tubule **(d)** and apparently fused cylindrical tubules **(e)**. Dashed lines indicate the approximate tubule boundaries.

**Extended Data Fig. 2.**
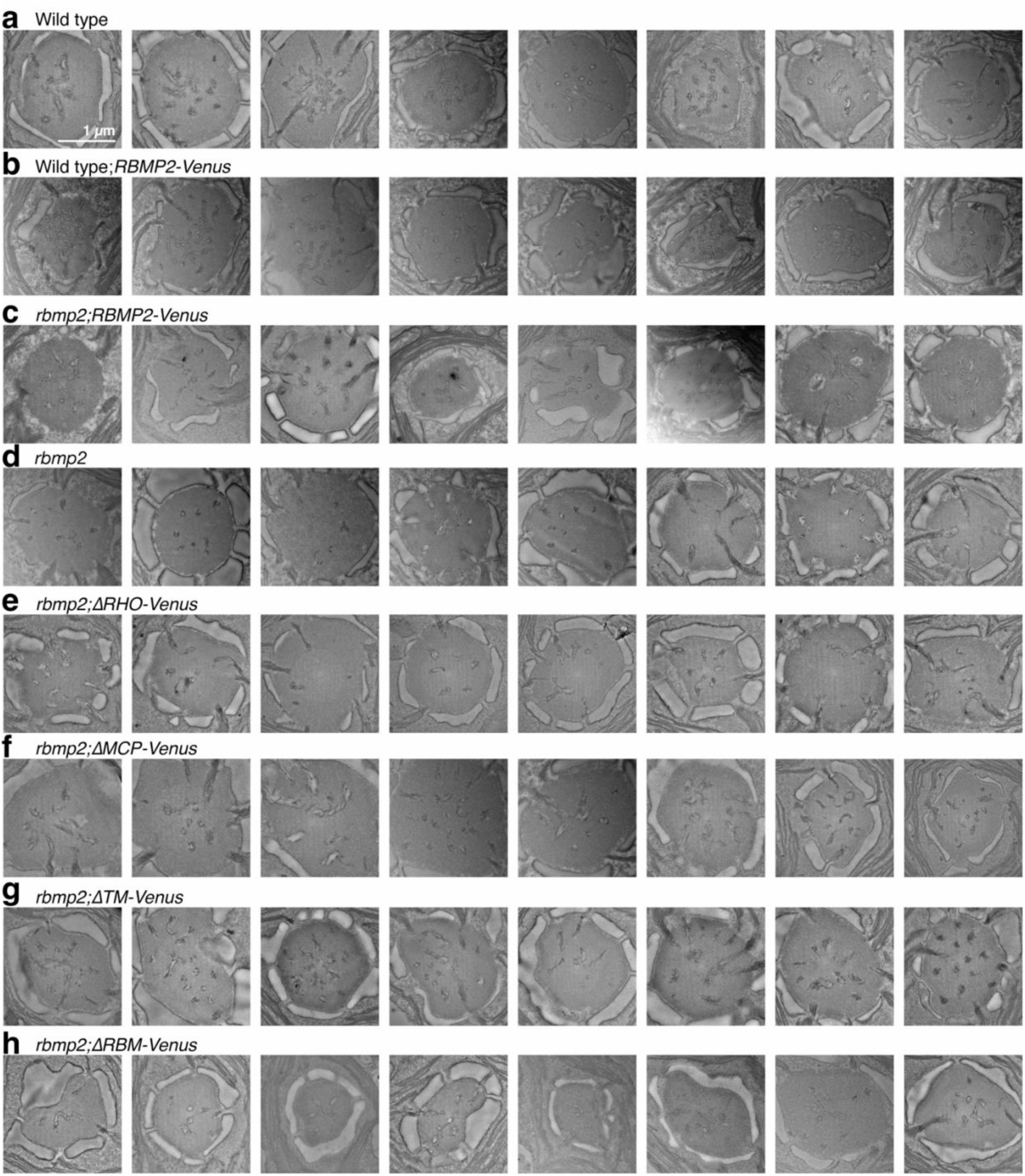
Additional TEM images of pyrenoids from wild type, wild-type;*RBMP2-Venus*, *rbmp2* and *rbmp2* strains expressing RBMP2 variants. **a–h,** Additional representative TEM images of pyrenoids from wild type (**a**), wild-type;*RBMP2-Venus* (**b**), *rbmp2;RBMP2-Venus* (**c**), *rbmp2* (**d**), *rbmp2;ΔRHO-Venus* (**e**), *rbmp2;ΔMCP-Venus* (**f**), *rbmp2;ΔTM-Venus* (**g**) and *rbmp2;ΔRBM-Venus* (**h**) cells. All images have the same scale.

**Extended Data Fig. 3.**
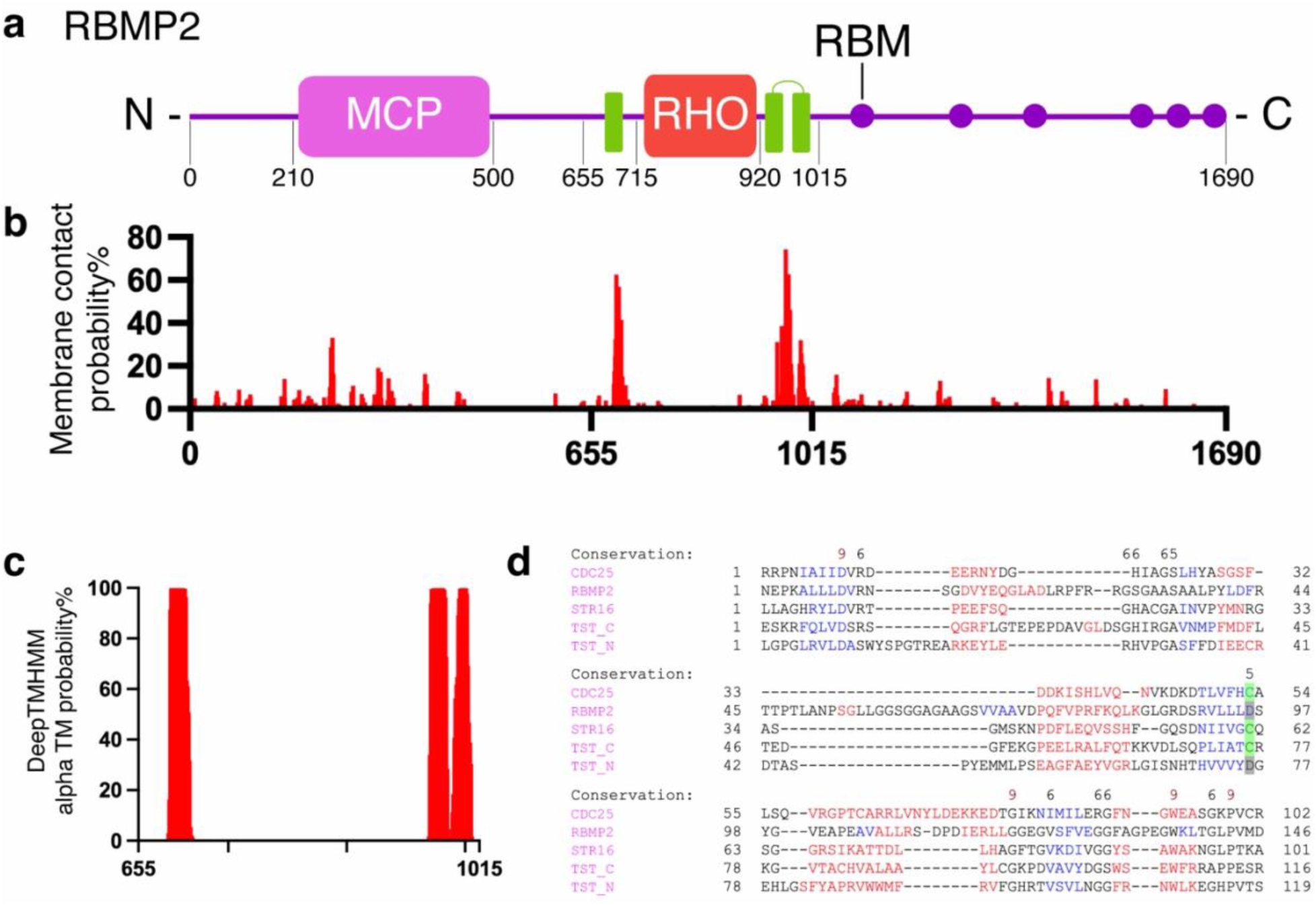
Domain analysis indicates that RBMP2 is a membrane-associated protein with a non-catalytic rhodanese domain. **a,** Schematic of RBMP2 showing predicted functional domains, including an N-terminal high membrane-contact-probability region, a central rhodanese domain, predicted transmembrane helices and six C-terminal Rubisco-binding motifs. **b,** Membrane-contact-probability scores for RBMP2 predicted by a sequence-based model that estimates the likelihood of individual residues contacting the lipid bilayer^33^. **c,** Predicted transmembrane helices in RBMP2 identified by DeepTMHMM^45^. **d,** Multiple sequence alignment of the RBMP2 rhodanese domain with representative rhodanese-family domains, including CDC25 and STR16 from *Arabidopsis thaliana*, and the N-terminal and C-terminal rhodanese domains of human thiosulfate sulfurtransferase (TST). The catalytic cysteine residue is marked in green, and the non-catalytic aspartic acid residue in RBMP2 is marked in grey.

**Extended Data Fig. 4.**
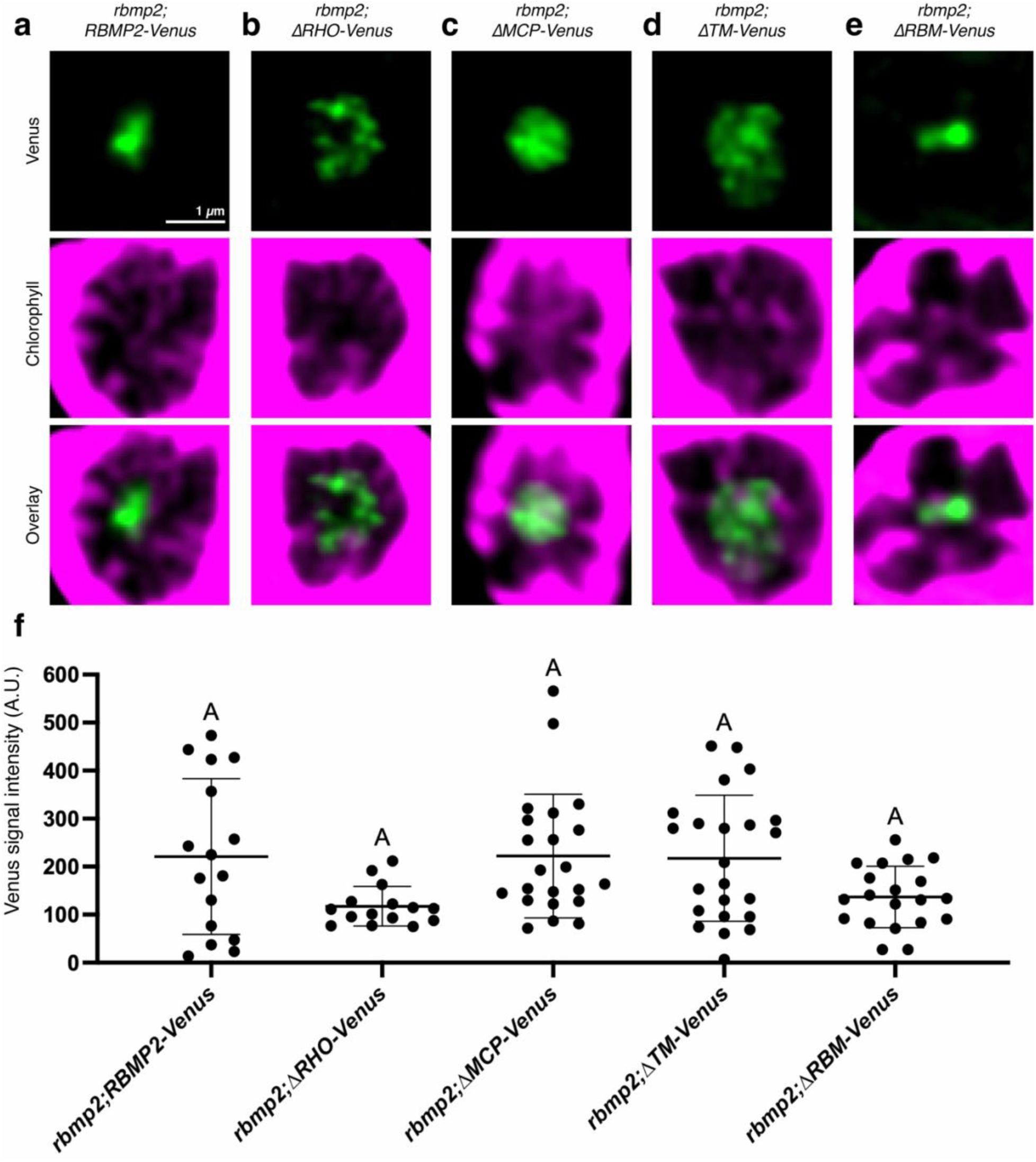
RBMP2 truncation variants localize to the pyrenoid and are expressed at similar levels. **a–e,** Live-cell confocal microscopy images showing the localization of Venus-tagged RBMP2 and RBMP2 truncation variants in the rbmp2 mutant background. Chlorophyll autofluorescence marks the chloroplast. **f,** Quantification of Venus signal intensity in the pyrenoid in *rbmp2;RBMP2-Venus*, *rbmp2;ΔRHO-Venus, rbmp2;ΔMCP-Venus*, *rbmp2;ΔTM-Venus* and *rbmp2;ΔRBM-Venus* cells. Background intensity measured from *rbmp2* mutant pyrenoids was subtracted from the measured intensity in Venus-expressing strains. Data were analyzed by one-way ANOVA with Tukey’s multiple-comparisons test. Error bars indicate s.d.

**Extended Data Fig. 5.**
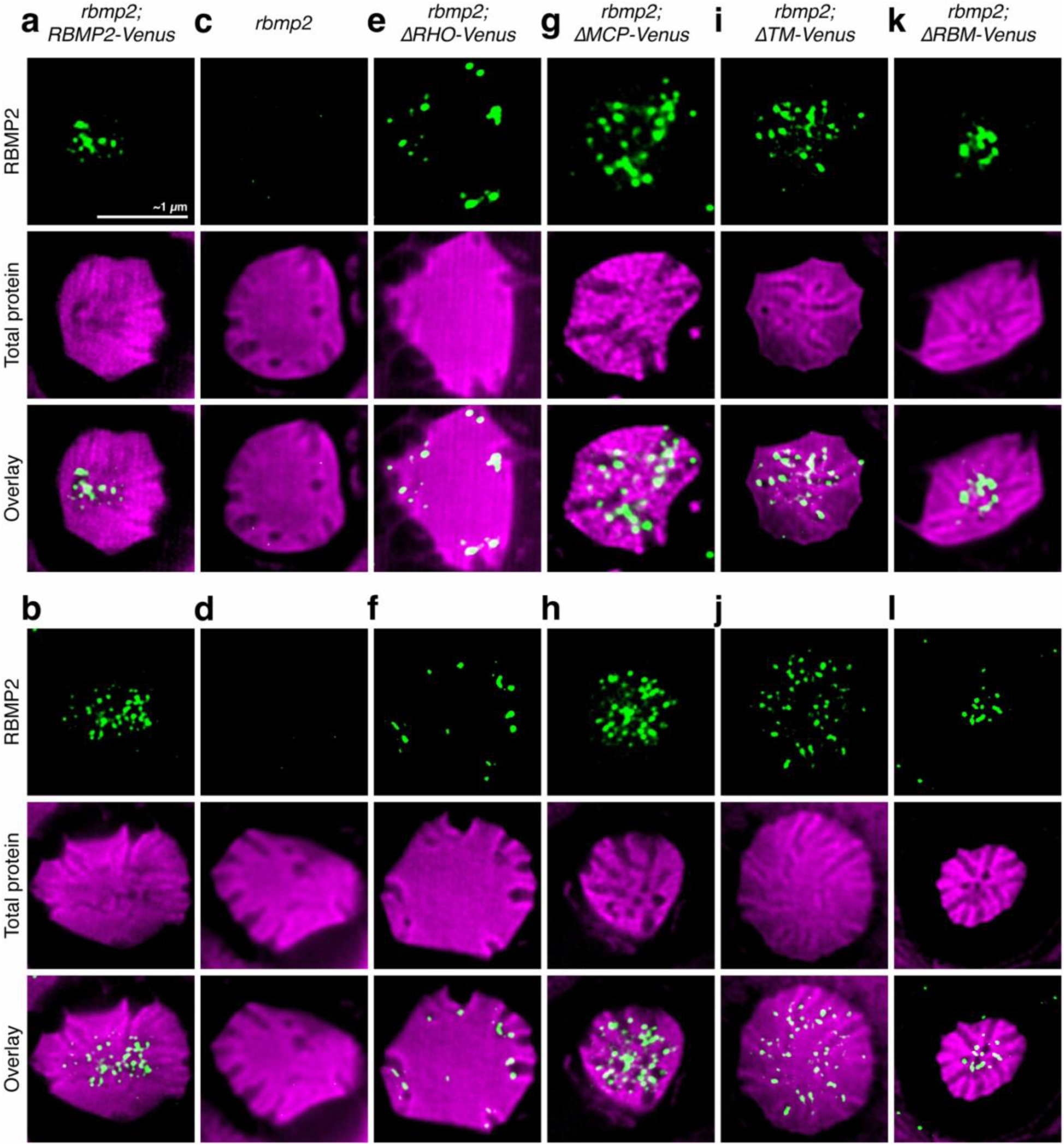
Additional ExM images of *rbmp2* strains expressing RBMP2 truncation variants. **a–l,** Additional ExM images of pyrenoids from *rbmp2;RBMP2-Venus* **(a,b)**, *rbmp2* **(c,d)**, *rbmp2;ΔRHO-Venus* **(e,f)**, *rbmp2;ΔMCP-Venus* **(g,h)**, *rbmp2;ΔTM-Venus* **(i,j)** and *rbmp2;ΔRBM-Venus* **(k,l)** cells. Samples were stained for FLAG-tagged RBMP2 variants by immunofluorescence and for total protein with NHS ester to visualize the pyrenoid matrix. Scale bars in ExM images are corrected for the expansion factor and represent estimated true cellular dimensions.

**Extended Data Fig. 6.**
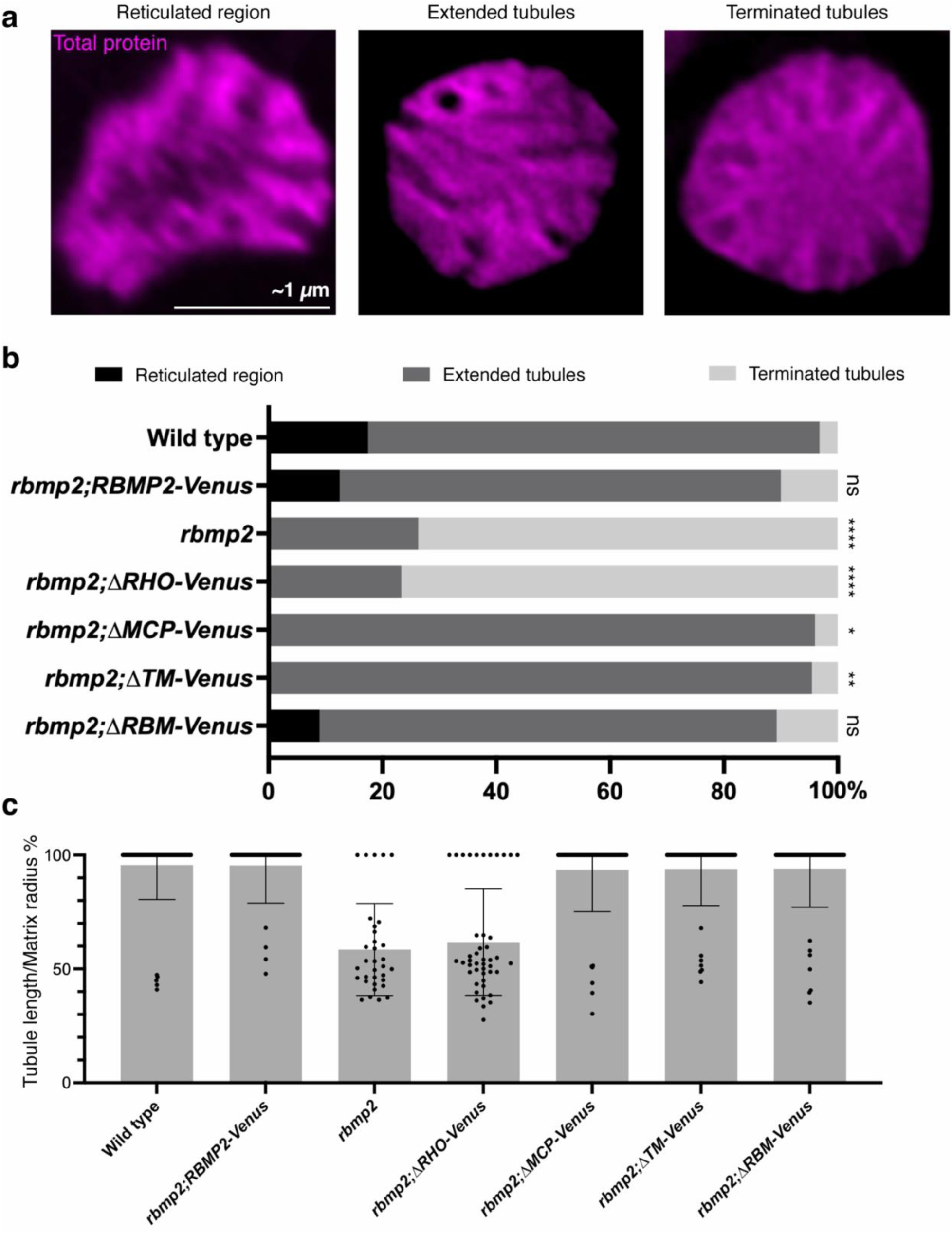
Tubule-network morphology differs among wild-type, *rbmp2* and RBMP2 truncation strains. **a,** Representative ExM images showing the three tubule-network phenotypes scored in this study: a reticulated region, extended tubules and terminated tubules. Representative images are from wild-type, *rbmp2;ΔTM-Venus* and *rbmp2* cells, respectively. Samples were stained for total protein with NHS ester to visualize the pyrenoid matrix. In some ExM images, connected tubule networks could not be confidently resolved as reticulated-region tubules or cylindrical tubules because of insufficient contrast. **b,** Percentage of cells showing each tubule-network phenotype in wild type, *rbmp2*, *rbmp2;RBMP2-Venus*, *rbmp2;ΔRHO-Venus*, *rbmp2;ΔMCP-Venus*, *rbmp2;ΔTM-Venus* and *rbmp2;ΔRBM-Venus* strains. Phenotype distributions for each strain were compared with wild type using Fisher’s exact test on 2 × 3 contingency tables. Number of cells scored: wild type, *n* = 63; *rbmp2;RBMP2-Venus*, *n* = 80; *rbmp2*, *n* = 38; *rbmp2;ΔRHO-Venus*, *n* = 30; *rbmp2;ΔMCP-Venus*, *n* = 29; *rbmp2;ΔTM-Venus*, *n* = 44; *rbmp2;ΔRBM-Venus*, *n* = 56. *=*P* < 0.05; **=*P* < 0.01; ****=*P* < 0.0001; ns, not significant (*P* > 0.05). **c,** Quantification of tubule extension, measured as tubule length divided by matrix radius, in wild-type, *rbmp2;RBMP2-Venus*, *rbmp2*, *rbmp2;ΔRHO-Venus*, *rbmp2;ΔMCP-Venus*, *rbmp2;ΔTM-Venus* and *rbmp2;ΔRBM-Venus* pyrenoids. Number of tubules measured: wild type, *n* = 63; *rbmp2;RBMP2-Venus*, *n* = 60; *rbmp2*, *n* = 33; *rbmp2;ΔRHO-Venus*, *n* = 45; *rbmp2;ΔMCP-Venus*, *n* = 51; *rbmp2;ΔTM-Venus*, *n* = 54; *rbmp2;ΔRBM-Venus*, *n* = 60. Error bars indicate s.d.

**Extended Data Fig. 7.**
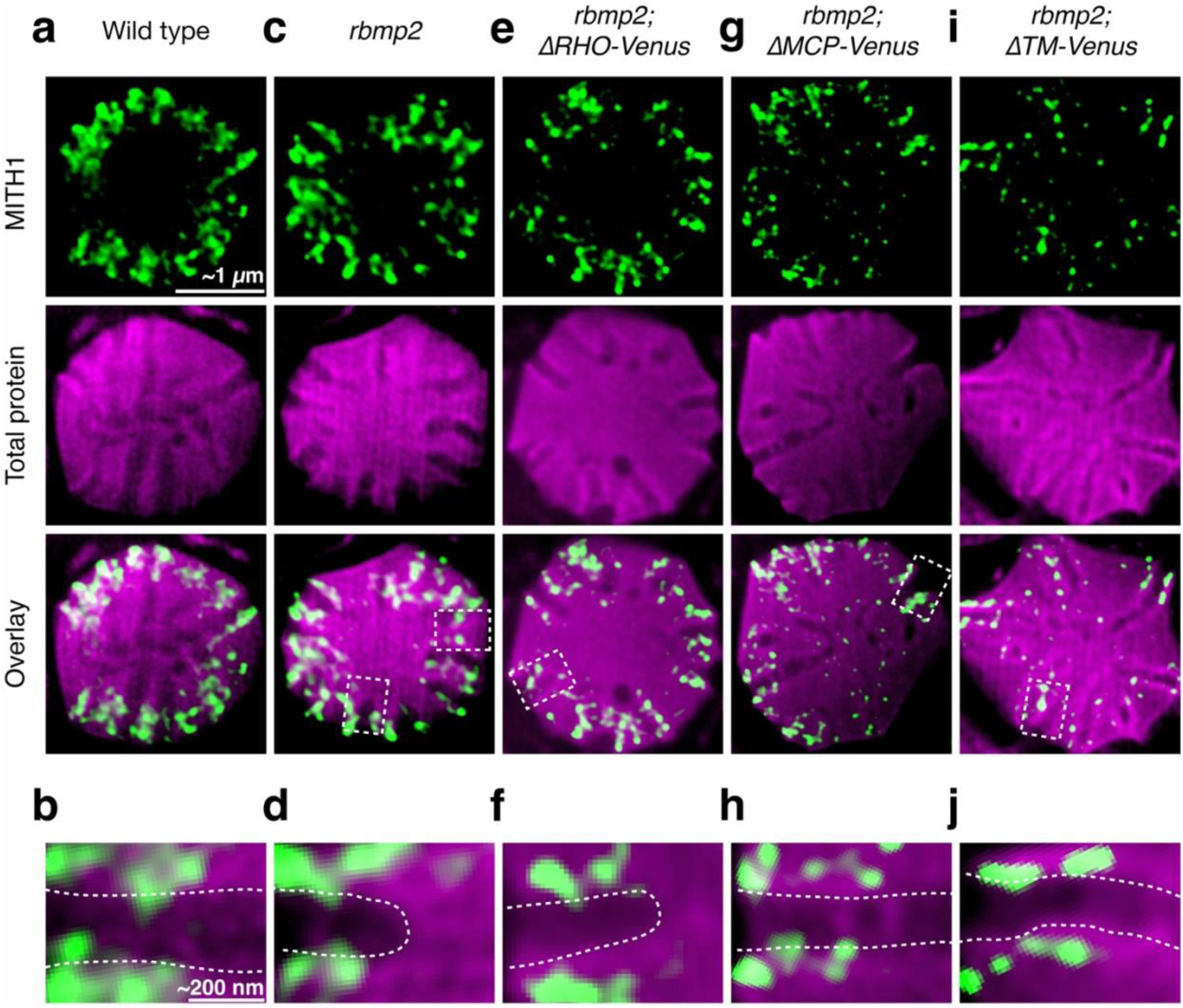
MITH1 primarily localizes to the peripheral portion of cylindrical tubules in RBMP2 variant strains. **a,c,e,g,i,** ExM images of wild-type, *rbmp2*, and *rbmp2* strains expressing the indicated RBMP2 variants. Samples were stained for MITH1 by immunofluorescence and for the total protein by NHS-ester staining. Images show MITH1, total protein, and merged channels. **b,d,f,h,j,** Magnified views of the boxed regions in **a,c,e,g,i**, respectively. Dashed lines indicate the approximate boundaries of cylindrical tubules.

**Extended Data Fig. 8.**
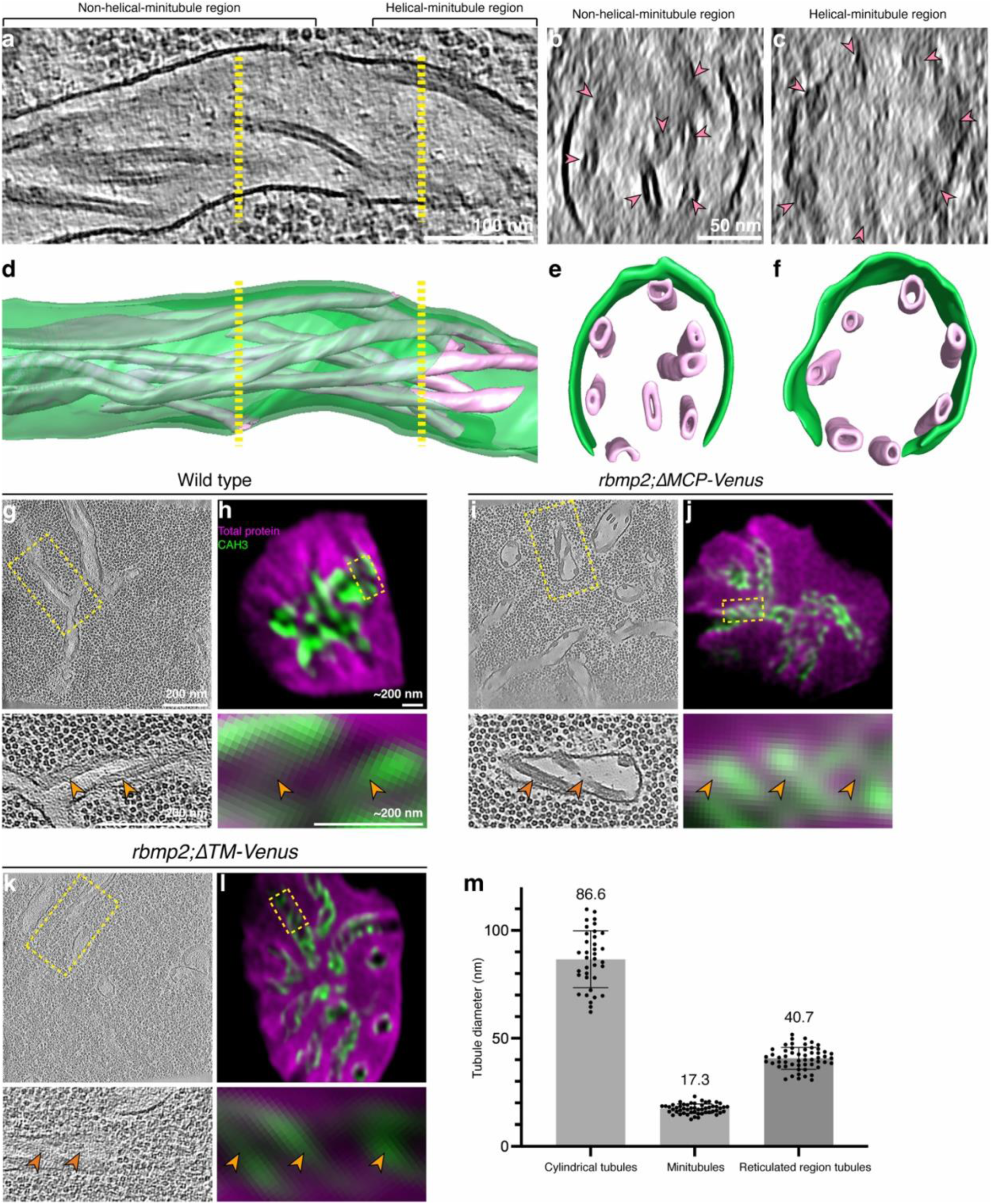
Minitubules transition from a non-helical-minitubule region to a helical-minitubule region as cylindrical tubules approach the pyrenoid center, and CAH3 localizes to stripes in wild-type and RBMP2 mutants. **a,** Slice from a tomogram of a wild-type *Chlamydomonas* pyrenoid showing a cylindrical tubule extending from the pyrenoid periphery (left) toward the pyrenoid center (right). Yellow dashed lines indicate the cross-sections shown in **b** and **c**. **b,c,** Cross-sections of the cylindrical tubule shown in **a**, showing the non-helical-minitubule region (**b**) and the helical-minitubule region (**c**). In the non-helical-minitubule region, some minitubules are positioned away from the cylindrical-tubule membrane; in the helical-minitubule region, all minitubules are closely apposed to the cylindrical-tubule membrane. Magenta arrowheads indicate minitubules. **d–f,** Segmentations of the tomogram and cross-sections shown in **a–c**, respectively. Cylindrical tubules are shown in green, and minitubules are shown in magenta. The segmentation in **f** shows helical minitubules along the luminal surface of the cylindrical tubule in the helical-minitubule region. **g–l,** Additional examples comparing stripe-like minitubule patterns observed by cryo-ET with CAH3 stripe patterns observed by ExM in wild-type (**g,h**), *rbmp2;ΔMCP-Venus* (**i,j**) and *rbmp2;ΔTM-Venus* (**k,l**) cells. In the cryo-ET images, yellow dashed boxes indicate the regions magnified below each tomogram slice. In the ExM images, cells were stained for CAH3 by immunofluorescence and for total protein with NHS ester; boxed regions are magnified below each image. Orange arrowheads indicate stripe-like minitubule patterns in cryo-ET and CAH3 stripes in ExM. **m,** Quantification of the diameters of cylindrical tubules, minitubules and reticulated-region tubules measured from cryo-ET data. Scale bars in ExM images are corrected for the expansion factor and represent estimated true cellular dimensions. Error bars indicate s.d.

**Extended Data Fig. 9.**
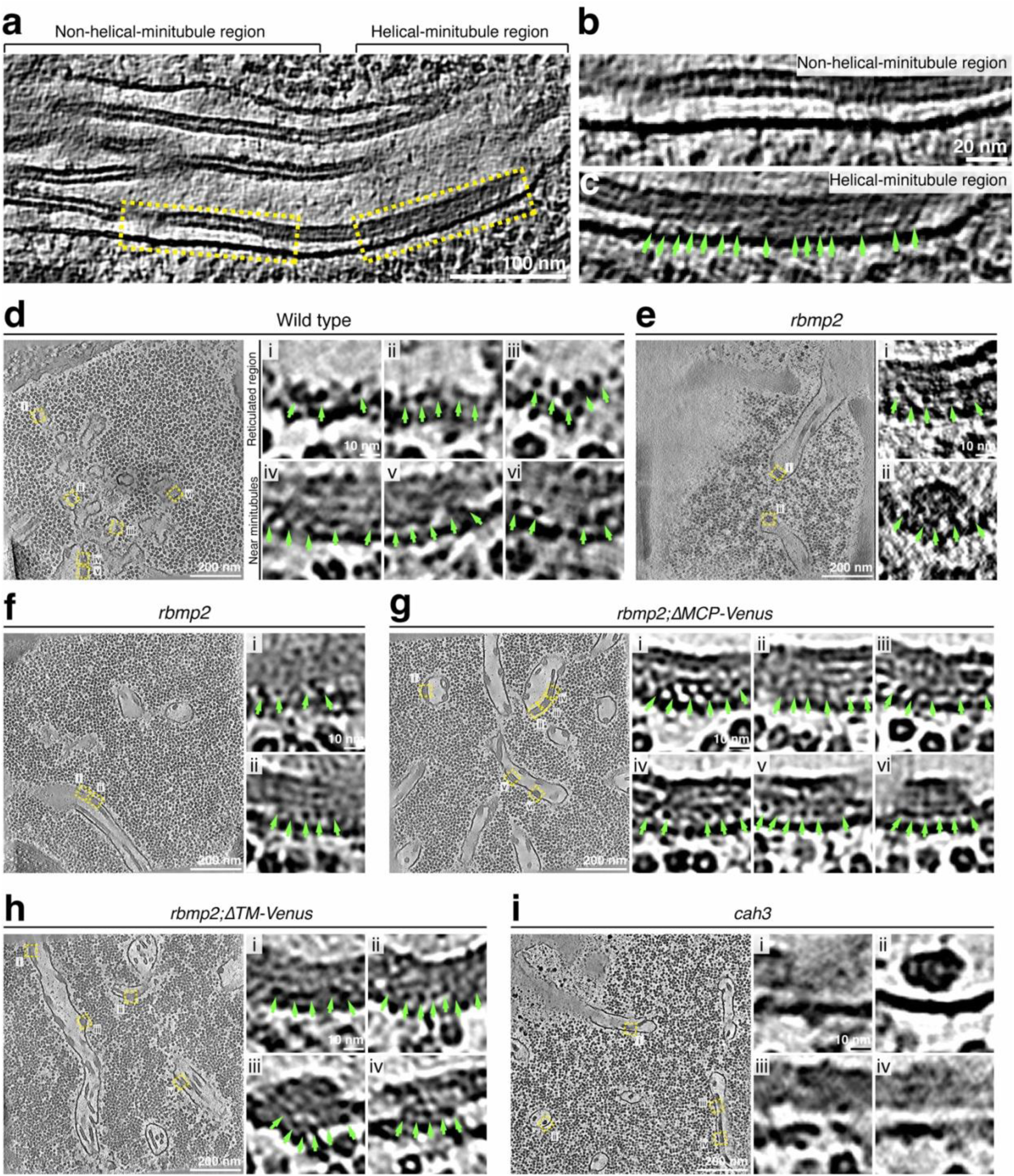
Additional examples of putative CAH3 densities on wild-type reticulated-region membranes and at the interface between the minitubules and cylindrical tubules in the helical-minitubule region across strains. **a,** Slice from a tomogram of a wild-type *Chlamydomonas* pyrenoid showing a cylindrical tubule extending from the pyrenoid periphery (left) toward the pyrenoid center (right). Yellow dashed lines indicate the magnified views in **b** and **c**. The slices shown in **a** were taken from the same tomogram as those shown in Extended Data Fig. 8,a,b,c. **b,c,** Magnified views of the boxed regions in **a**, showing the non-helical-minitubule region (**b**) and the helical-minitubule region (**c**). Green arrowheads indicate densities observed near minitubules in the helical-minitubule region. The slices shown in **b,c** were taken from the same tomogram as those shown in Extended Data Fig. 8,a,b,c. **d,** Additional tomogram slices and magnified views showing densities in wild-type cells (CC-124). Panels i–iii show densities on the reticulated-region membrane, and panels iv–vi show densities at the interface between minitubules and the cylindrical-tubule membrane. Yellow dashed boxes indicate regions shown in the corresponding magnified views. Green arrowheads indicate the densities. **e–h,** Additional tomogram slices and magnified views showing densities at the interface between minitubules and the cylindrical-tubule membrane in the helical-minitubule region in *rbmp2* cells (**e,f**), *rbmp2;ΔMCP-Venus* (**g**) and *rbmp2;ΔTM-Venus* (**h**) cells. Yellow dashed boxes indicate regions shown in the corresponding magnified views. The slices shown in **e** were taken from the same tomogram as those shown in Fig. 3e,g and Fig. 5o,p; those shown in **g** were taken from the same tomogram as that shown in Extended Data Fig. 8,i; and those shown in **h** were taken from the same tomogram as those shown in Fig. 5i,s,t. Green arrowheads indicate densities at the interface between minitubules and the cylindrical-tubule membrane. **i,** Additional tomogram slices and magnified views showing the absence of densities at the interface between minitubules and the cylindrical-tubule membrane in *cah3* cells. The slices shown in **i** were taken from the same tomogram as those shown in Fig. 5u,v.

**Extended Data Fig. 10.**
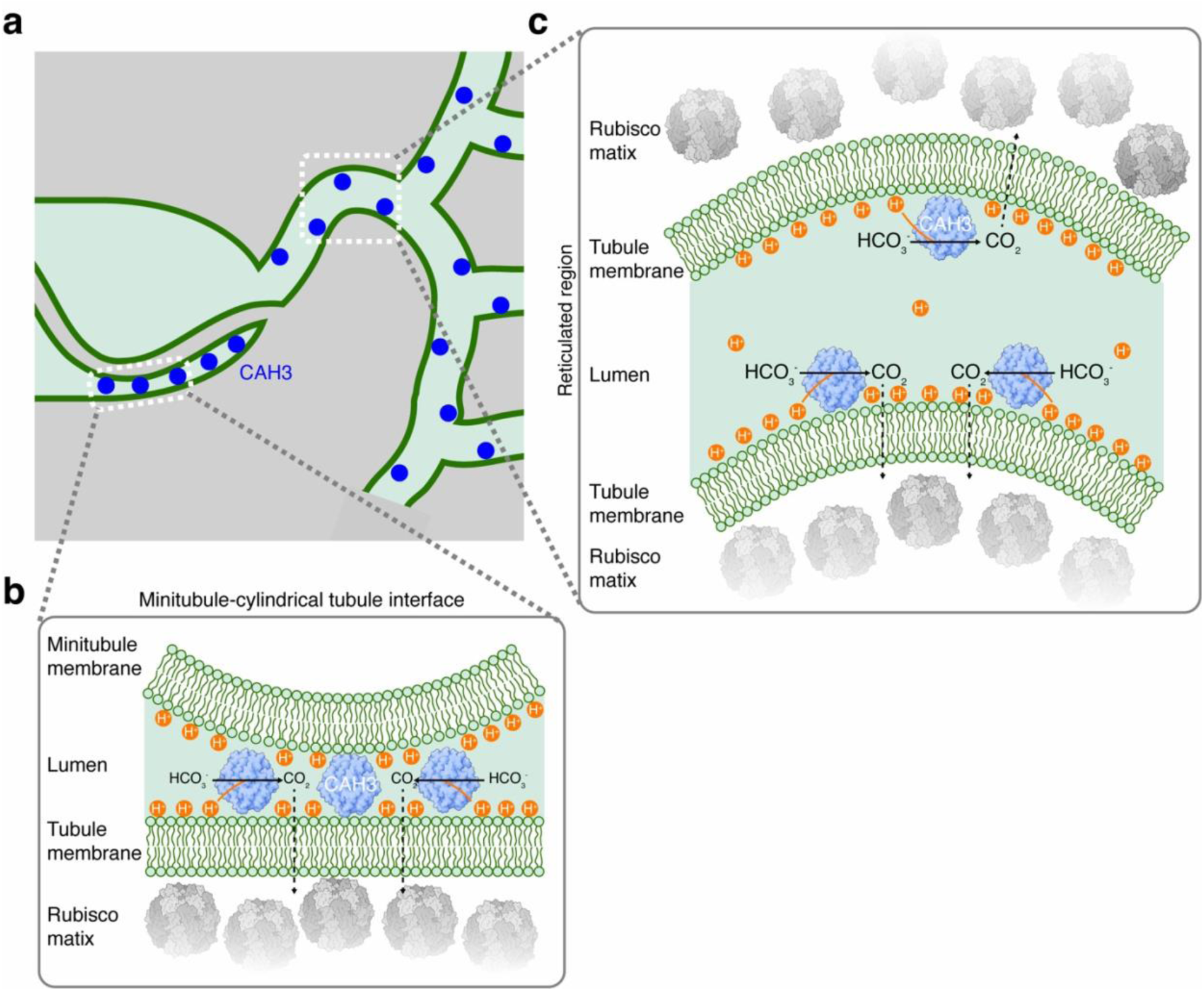
Speculative model for CAH3 function at pyrenoid membranes. **a,** Model for CAH3 localization in wild-type pyrenoids. CAH3 localizes to the reticulated region and to the interface between minitubules and cylindrical tubules. **b,** Speculative model for proton distribution at the minitubule–cylindrical tubule interface. **c,** Speculative model for proton distribution in the reticulated region.

**Supplementary Fig. 1.**
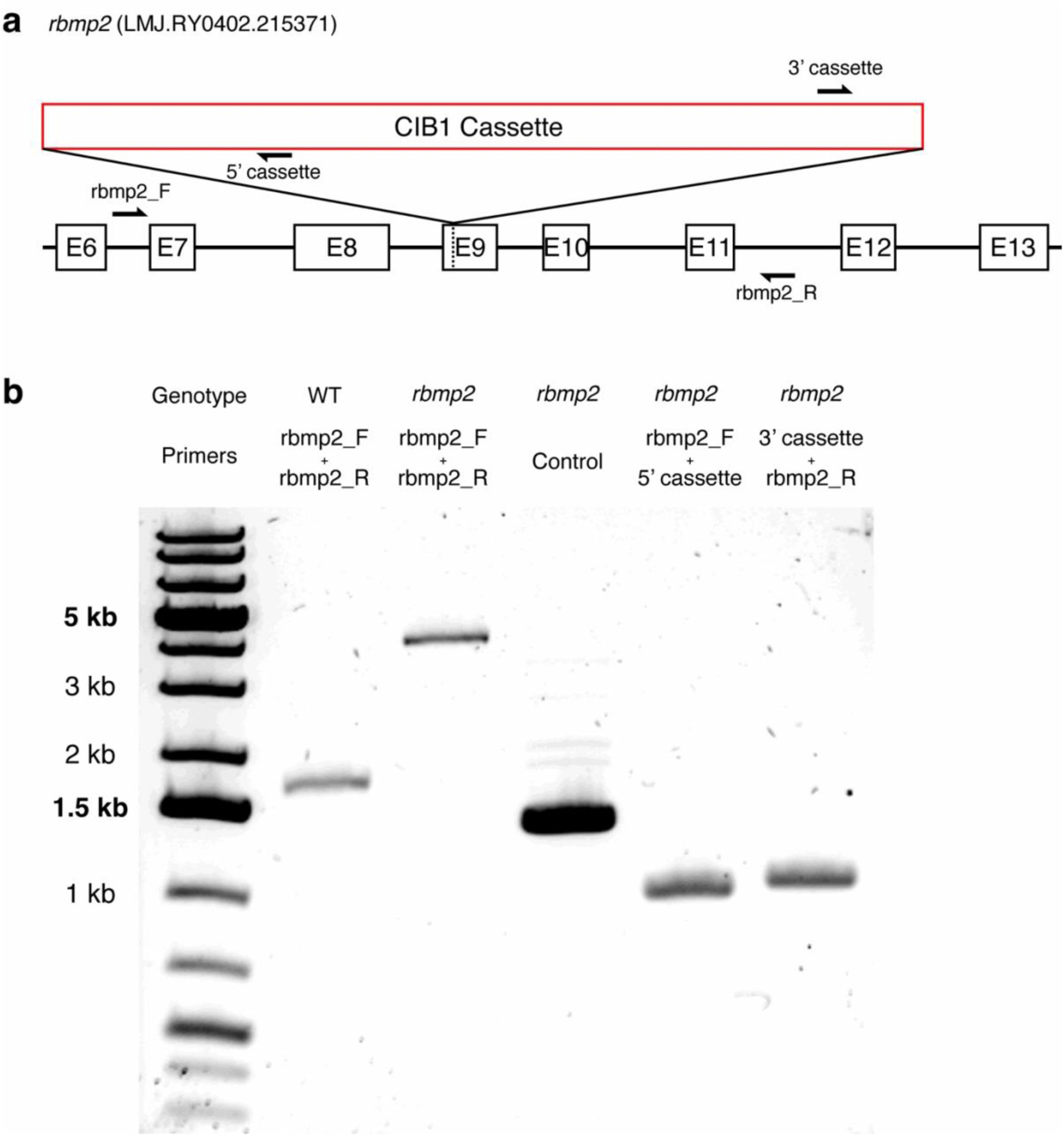
PCR genotyping confirms the insertion in the rbmp2 mutant. **a,** Schematic of the RBMP2 locus in the *rbmp2* mutant strain LMJ.RY0402.215371, showing the position of the CIB1 insertion cassette and the primers used for PCR genotyping. The CIB1 cassette is 2,223 bp. The exon primer pair *rbmp2_F/rbmp2_R* is expected to produce a 1,655-bp product from wild-type genomic DNA. The 5′ cassette primer pair *rbmp2_F/5′ cassette* is expected to produce a 939-bp product from *rbmp2* genomic DNA, and the 3′ cassette primer pair *3′ cassette/rbmp2_R* is expected to produce a 973-bp product from *rbmp2* genomic DNA. **b,** Agarose gel showing PCR amplification from genomic DNA isolated from wild-type and *rbmp2* cells. Amplification with rbmp2_F/rbmp2_R produced the expected wild-type product in wild type and a larger product in *rbmp2*, consistent with cassette insertion. PCRs using cassette-specific primers amplified the insertion junctions in *rbmp2*. Control primers were used to confirm genomic DNA quality.

**Supplementary Fig. 2.**
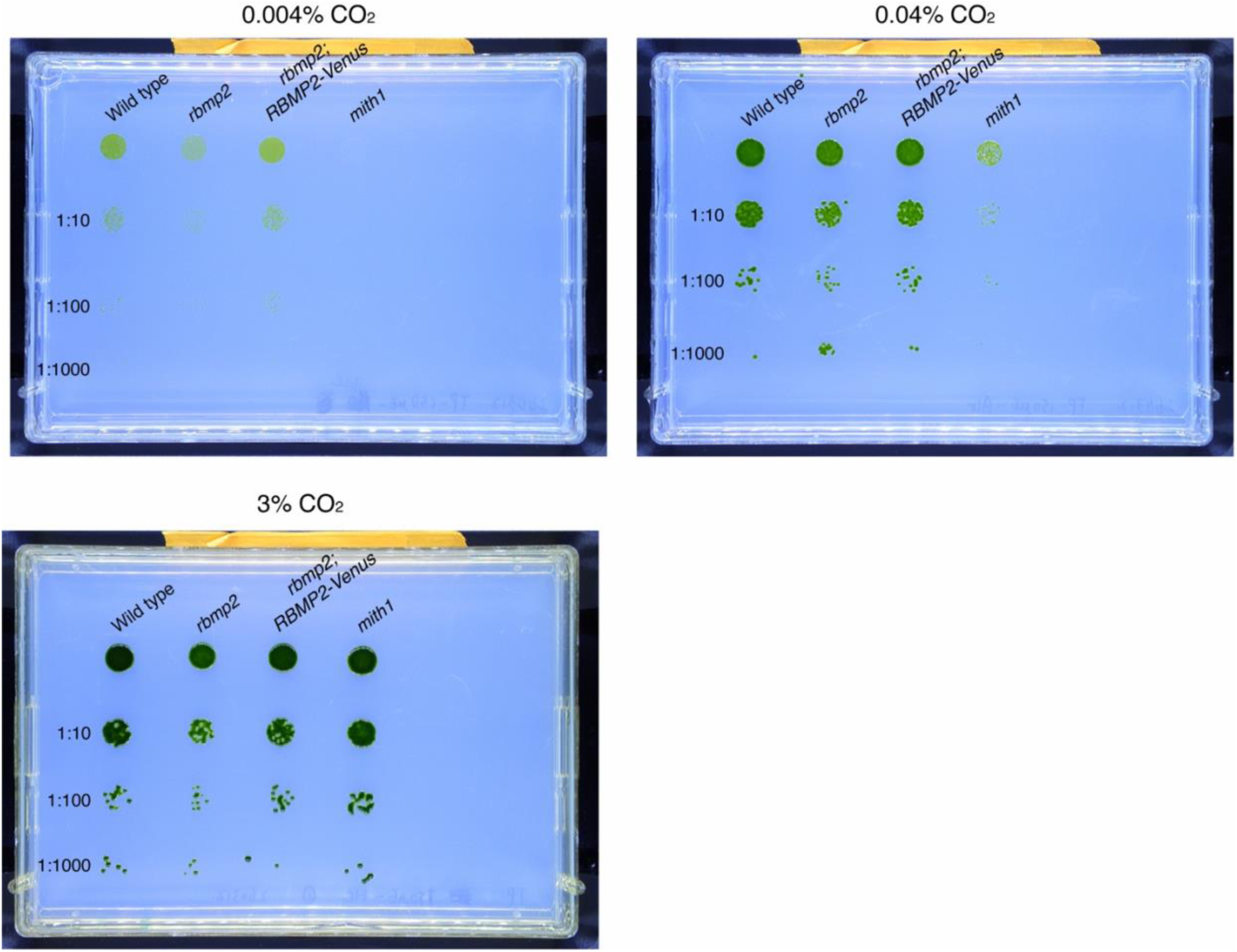
Full plate images for the growth assay. Full plate images corresponding to the spot growth assay shown in Fig. 1e. Cells were spotted on agar medium at the indicated serial dilutions and grown under very low CO₂, air-level CO₂, or high CO₂ conditions. Spot dilutions were 1:1, 1:10, 1:100 and 1:1000.

**Supplementary Fig. 3.**
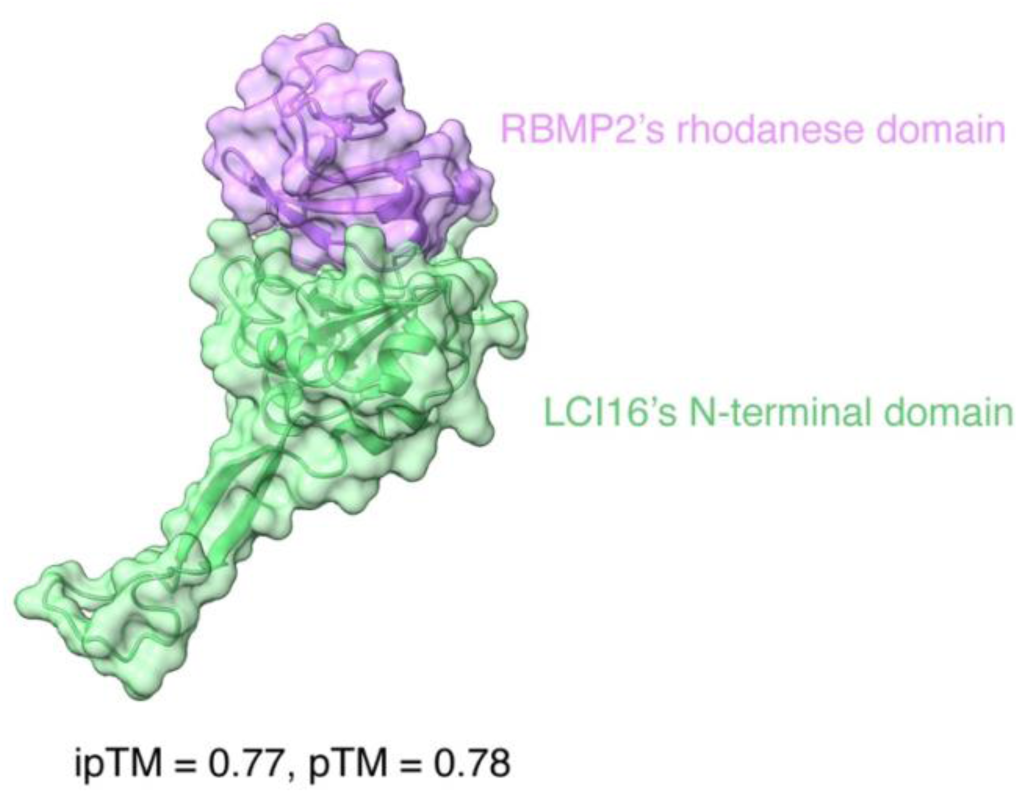
AlphaFold2 predicts an interaction between LCI16 and the RBMP2 rhodanese domain. AlphaFold2 prediction of the N-terminal domain of LCI16 bound to the rhodanese domain of RBMP2. The predicted complex is shown as a surface and cartoon representation, with LCI16 in green and the RBMP2 rhodanese domain in purple. The prediction had an ipTM score of 0.77 and a pTM score of 0.78.

**Supplementary Table 1.**
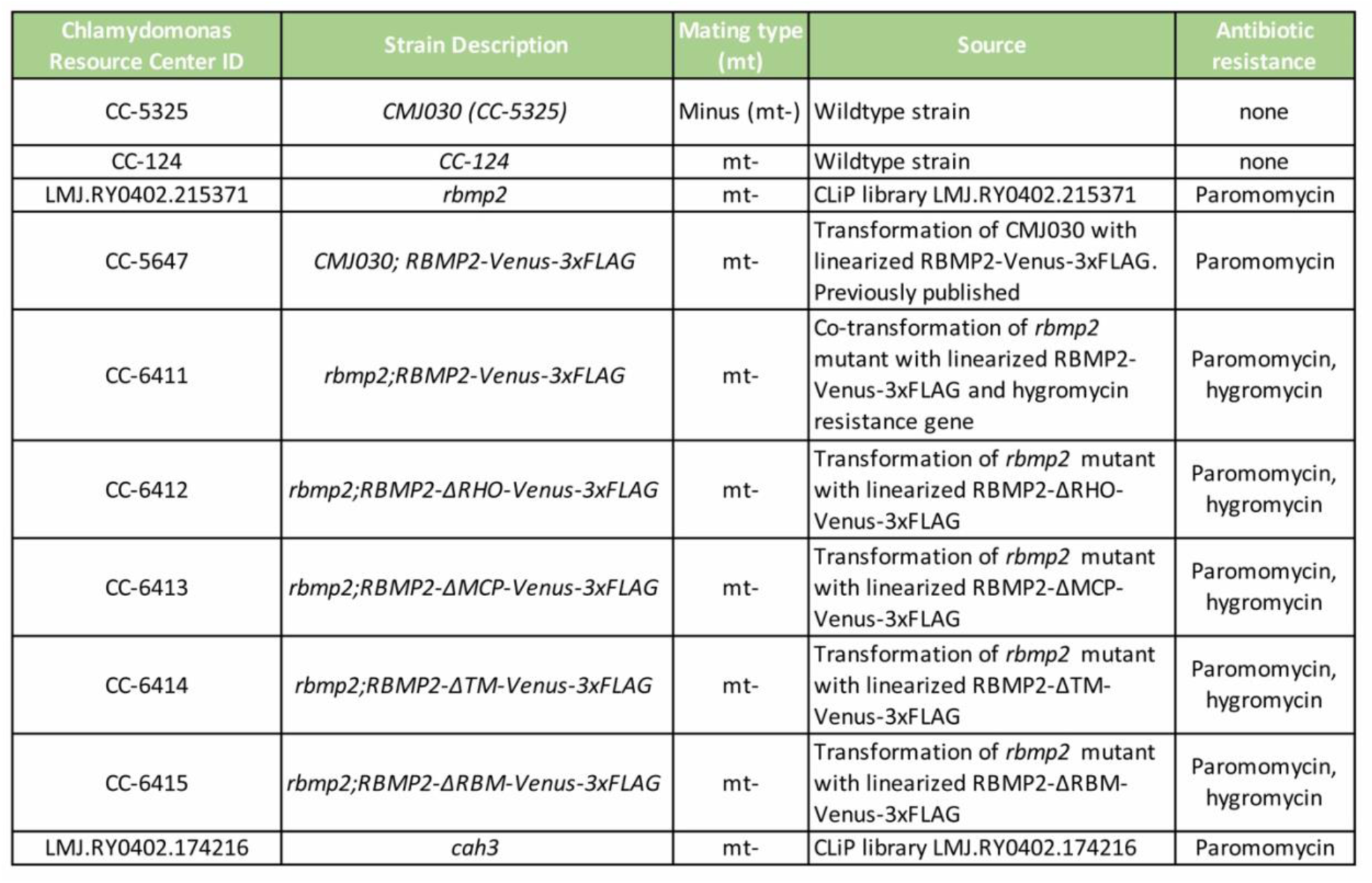
Chlamydomonas strains used in this study. List of Chlamydomonas strains used in this study and their sources. CMJ030 is the wild-type background strain used for the CLiP mutant library. The *rbmp2* and *cah3* mutants were generated in CLiP mutant library study through insertional mutagenesis with a paromomycin-resistance cassette and were obtained from the CLiP library^46^. CC-5647 and CC-124 were described previously^27,47^. New strains generated in this study were produced by transformation.

| Chlamydomonas Resource Center ID | Strain Description | Mating type (mt) | Source | Antibiotic resistance |
| --- | --- | --- | --- | --- |
| CC-5325 | *CMJ030 (CC-5325)* | Minus (mt-) | Wildtype strain | none |
| CC-124 | *CC-124* | mt- | Wildtype strain | none |
| LMJ.RY0402.215371 | *rbmp2* | mt- | CLiP library LMJ.RY0402.215371 | Paromomycin |
| CC-5647 | *CMJ030; RBMP2-Venus-3xFLAG* | mt- | Transformation of *CMJ030* with linearized *RBMP2-Venus-3xFLAG*. Previously published | Paromomycin |
| CC-6411 | *rbmp2;RBMP2-Venus-3xFLAG* | mt- | Co-transformation of *rbmp2* mutant with linearized *RBMP2-Venus-3xFLAG* and hygromycin resistance gene | Paromomycin, hygromycin |
| CC-6412 | *rbmp2;RBMP2-ΔRHO-Venus-3xFLAG* | mt- | Transformation of *rbmp2* mutant with linearized *RBMP2-ΔRHO-Venus-3xFLAG* | Paromomycin, hygromycin |
| CC-6413 | *rbmp2;RBMP2-ΔMCP-Venus-3xFLAG* | mt- | Transformation of *rbmp2* mutant with linearized *RBMP2-ΔMCP-Venus-3xFLAG* | Paromomycin, hygromycin |
| CC-6414 | *rbmp2;RBMP2-ΔTM-Venus-3xFLAG* | mt- | Transformation of *rbmp2* mutant with linearized *RBMP2-ΔTM-Venus-3xFLAG* | Paromomycin, hygromycin |
| CC-6415 | *rbmp2;RBMP2-ΔRBM-Venus-3xFLAG* | mt- | Transformation of *rbmp2* mutant with linearized *RBMP2-ΔRBM-Venus-3xFLAG* | Paromomycin, hygromycin |
| LMJ.RY0402.174216 | *cah3* | mt- | CLiP library LMJ.RY0402.174216 | Paromomycin |

**Supplementary Table 2.** Plasmids used for Chlamydomonas transformation. List of plasmids used for transformation into Chlamydomonas. The restriction enzyme used to linearize each plasmid before transformation is indicated. The pLM155 plasmid was generated and described in a previous study^27^.

| Plasmid | Source | *Chlamydomonas*<br>antibiotic<br>resistance | *E. Coli*<br>antibiotic<br>resistance | Restriction<br>enzymes |
| --- | --- | --- | --- | --- |
| RBMP2-Venus-3xFLAG,<br>pLM155 | Previously published | Paromomycin | Kanamycin | XbaI |
| RBMP2-ΔRHO-Venus-3xFLAG, pHW013 | Commercially synthesized RBMP2<br>lacking the rhodanese domain from<br>pLM155 and switched with the Ori<br>and antibiotic resistance genes from<br>pRAM118 | Hygromycin | Ampicillin | NdeI |
| RBMP2-ΔMCP-Venus-3xFLAG, pHW008 | Commercially synthesized RBMP2<br>lacking the MCP domain from<br>pLM155 and switched with the Ori<br>and antibiotic resistance genes from<br>pRAM118 | Hygromycin | Ampicillin | NdeI |
| RBMP2-ΔTM-Venus-3xFLAG, pHW012 | Commercially synthesized RBMP2<br>lacking the transmembrane domains<br>from pLM155 and switched with the<br>Ori and antibiotic resistance genes<br>from pRAM118 | Hygromycin | Ampicillin | NdeI |
| RBMP2-ΔRBM-Venus-3xFLAG, pHW010 | Commercially synthesized RBMP2<br>lacking 6 Rubisco-binding motifs<br>from pLM155 and switched with the<br>Ori and antibiotic resistance genes<br>from pRAM118 | Hygromycin | Ampicillin | NdeI |

## REFERENCES

1. Field, C. B., Behrenfeld, M. J., Randerson, J. T. & Falkowski, P. Primary Production of the Biosphere: Integrating Terrestrial and Oceanic Components. Science 281, 237–240 (1998).

2. Mackinder, L. C. M. et al. A repeat protein links Rubisco to form the eukaryotic carbon-concentrating organelle. Proceedings of the National Academy of Sciences 113, 5958–5963 (2016).

3. Behrenfeld, M. J. et al. Biospheric Primary Production During an ENSO Transition. Science 291, 2594–2597 (2001).

4. Rousseaux, C. S. & Gregg, W. W. Interannual Variation in Phytoplankton Primary Production at A Global Scale. Remote Sensing 6, 1–19 (2014).

5. G, M. D. Chloroplast morphology, movements and inheritance in diatoms. Cytology, Genetics and Molecular Biology of Algae https://cir.nii.ac.jp/crid/1571698599039433728 (1996).

6. Thierstein, H. R. & Young, J. R. Coccolithophores: From Molecular Processes to Global Impact. (Springer Science & Business Media, 2013).

7. Meyer, M. & Griffiths, H. Origins and diversity of eukaryotic CO2-concentrating mechanisms: lessons for the future. J Exp Bot 64, 769–786 (2013).

8. Rochaix, J.-D. The Pyrenoid: An Overlooked Organelle Comes out of Age. Cell 171, 28–29 (2017).

9. Goodenough, U. W. Chloroplast Division and Pyrenoid Formation in Chlamydomonas Reinhardi. Journal of Phycology 6, 1–6 (1970).

10. Freeman Rosenzweig, E. S., et al. The Eukaryotic CO2-Concentrating Organelle Is Liquid-like and Exhibits Dynamic Reorganization. Cell 171, 148–162.e19 (2017).

11. He, S., Crans, V. L. & Jonikas, M. C. The pyrenoid: the eukaryotic CO2-concentrating organelle. Plant Cell 35, 3236–3259 (2023).

12. Barrett, J., Nam, O., Naduthodi, M. I. S. & Mackinder, L. C. M. Pyrenoid Structure, Function, Evolution, and Characterization Across Diverse Lineages. 10.1146/annurev-arplant-070225-034846 (2026) doi:10.1146/annurev-arplant-070225-034846.

13. Catherall, E. et al. From algae to plants: understanding pyrenoid-based CO2-concentrating mechanisms. Trends in Biochemical Sciences 50, 33–45 (2025).

14. He, S. et al. Kinase KEY1 controls pyrenoid condensate size throughout the cell cycle by disrupting phase separation interactions. Nat Cell Biol 28, 725–738 (2026).

15. Barrett, J., Girr, P. & Mackinder, L. C. M. Pyrenoids: CO2-fixing phase separated liquid organelles. Biochimica et Biophysica Acta (BBA) - Molecular Cell Research 1868, 118949 (2021).

16. Engel, B. D. et al. Native architecture of the Chlamydomonas chloroplast revealed by in situ cryo-electron tomography. eLife 4, e04889 (2015).

17. Itakura, A. K. et al. A Rubisco-binding protein is required for normal pyrenoid number and starch sheath morphology in Chlamydomonas reinhardtii. Proceedings of the National Academy of Sciences 116, 18445–18454 (2019).

18. Hennacy, J. H. et al. SAGA1 and MITH1 produce matrix-traversing membranes in the CO2-fixing pyrenoid. Nat. Plants 10, 2038–2051 (2024).

19. Garde, A., Wu, H. & Jonikas, M. C. Super-resolution expansion microscopy reveals nanoscale protein domains and CO2-dependent remodeling of Chlamydomonas pyrenoid-traversing membranes. Preprint at 10.64898/2026.06.11.731689 (2026).

20. Semenenko, V. E., P., N. A. Role of the pyrenoid in concentration, generation, and fixation of CO2 in the chloroplast of microalgae. Sov. Plant Physiol. 39, 470–476 (1992).

21. Raven, J. A. CO2-concentrating mechanisms: a direct role for thylakoid lumen acidification? Plant, Cell & Environment 20, 147–154 (1997).

22. Karlsson, J. et al. A novel alpha-type carbonic anhydrase associated with the thylakoid membrane in Chlamydomonas reinhardtii is required for growth at ambient CO2. EMBO J 17, 1208–1216 (1998).

23. Adler, L. et al. New horizons for building pyrenoid-based CO2-concentrating mechanisms in plants to improve yields. Plant Physiol 190, 1609–1627 (2022).

24. Fei, C., Wilson, A. T., Mangan, N. M., Wingreen, N. S. & Jonikas, M. C. Modelling the pyrenoid-based CO2-concentrating mechanism provides insights into its operating principles and a roadmap for its engineering into crops. Nat. Plants 8, 583–595 (2022).

25. Hennacy, J. H. & Jonikas, M. C. Prospects for Engineering Biophysical CO2 Concentrating Mechanisms into Land Plants to Enhance Yields. Annual Review of Plant Biology 71, 461–485 (2020).

26. Snead, W. T. & Gladfelter, A. S. The Control Centers of Biomolecular Phase Separation: How Membrane Surfaces, PTMs, and Active Processes Regulate Condensation. Molecular Cell 76, 295–305 (2019).

27. Meyer, M. T. et al. Assembly of the algal CO2-fixing organelle, the pyrenoid, is guided by a Rubisco-binding motif. Science Advances 6, eabd2408 (2020).

28. Yu, C.-C. (Jay) et al. Expansion microscopy of C. elegans. eLife 9, e46249 (2020).

29. Mao, C. et al. Feature-rich covalent stains for super-resolution and cleared tissue fluorescence microscopy. Science Advances 6, eaba4542 (2020).

30. M’Saad, O. & Bewersdorf, J. Light microscopy of proteins in their ultrastructural context. Nat Commun 11, 3850 (2020).

31. Dobro, M. J. et al. Uncharacterized Bacterial Structures Revealed by Electron Cryotomography. Journal of Bacteriology 199, 10.1128/jb.00100-17 (2017).

32. Bieber, A. et al. In situ structural analysis reveals membrane shape transitions during autophagosome formation. Proceedings of the National Academy of Sciences 119, e2209823119 (2022).

33. Wang, L., Zhang, J., Wang, D. & Song, C. Membrane contact probability: An essential and predictive character for the structural and functional studies of membrane proteins. PLOS Computational Biology 18, e1009972 (2022).

34. Ergun, S. L. et al. A bifunctional coiled-coil protein generates the membrane-within-condensate architecture of the CO2-fixing pyrenoid. 2026.06.09.731149 Preprint at 10.64898/2026.06.09.731149 (2026).

35. Wollweber, F. et al. Microtubules in Asgard archaea. Cell 188, 2451–2464.e26 (2025).

36. Kumar, G. et al. Sticker number modulates pyrenoid condensate assembly to support algal fitness. 2026.01.27.701992 Preprint at 10.64898/2026.01.27.701992 (2026).

37. Bordo, D. & Bork, P. The rhodanese/Cdc25 phosphatase superfamily. EMBO reports 3, 741–746 (2002).

38. Franklin, E. et al. Proteomic analysis of the pyrenoid-traversing membranes of Chlamydomonas reinhardtii reveals novel components. New Phytologist 249, 359–372 (2026).

39. Jarsch, I. K., Daste, F. & Gallop, J. L. Membrane curvature in cell biology: An integration of molecular mechanisms. J Cell Biol 214, 375–387 (2016).

40. Long, B. M., Förster, B., Pulsford, S. B., Price, G. D. & Badger, M. R. Rubisco proton production can drive the elevation of CO2 within condensates and carboxysomes. Proceedings of the National Academy of Sciences 118, e2014406118 (2021).

41. Yamano, T., Sato, E., Iguchi, H., Fukuda, Y. & Fukuzawa, H. Characterization of cooperative bicarbonate uptake into chloroplast stroma in the green alga Chlamydomonas reinhardtii. Proceedings of the National Academy of Sciences 112, 7315–7320 (2015).

42. Heberle, J., Riesle, J., Thiedemann, G., Oesterhelt, D. & Dencher, N. A. Proton migration along the membrane surface and retarded surface to bulk transfer. Nature 370, 379–382 (1994).

43. Springer, A., Hagen, V., Cherepanov, D. A., Antonenko, Y. N. & Pohl, P. Protons migrate along interfacial water without significant contributions from jumps between ionizable groups on the membrane surface. Proceedings of the National Academy of Sciences 108, 14461–14466 (2011).

44. Gennis, R. B. Proton Dynamics at the Membrane Surface. Biophysical Journal 110, 1909–1911 (2016).

45. Hallgren, J. et al. DeepTMHMM predicts alpha and beta transmembrane proteins using deep neural network. Preprint at 10.1101/2022.04.08.487609 (2022).

46. Li, X. et al. A genome-wide algal mutant library and functional screen identifies genes required for eukaryotic photosynthesis. Nat Genet 51, 627–635 (2019).

47. Pröschold, T., Harris, E. H. & Coleman, A. W. Portrait of a Species: Chlamydomonas reinhardtii. Genetics 170, 1601–1610 (2005).

48. Gambarotto, D. et al. Imaging cellular ultrastructures using expansion microscopy (U-ExM). Nat Methods 16, 71–74 (2019).

49. Damstra, H. G. et al. Visualizing cellular and tissue ultrastructure using Ten-fold Robust Expansion Microscopy (TREx). eLife 11, e73775 (2022).

50. Medeiros, J. M. et al. Robust workflow and instrumentation for cryo-focused ion beam milling of samples for electron cryotomography. Ultramicroscopy 190, 1–11 (2018).

51. Mastronarde, D. N. Automated electron microscope tomography using robust prediction of specimen movements. Journal of Structural Biology 152, 36–51 (2005).

52. Zheng, S. Q. et al. MotionCor2: anisotropic correction of beam-induced motion for improved cryo-electron microscopy. Nat Methods 14, 331–332 (2017).

53. Burt, A. et al. An image processing pipeline for electron cryo-tomography in RELION-5. FEBS Open Bio 14, 1788–1804 (2024).

54. Zheng, S., et al. AreTomo: An integrated software package for automated marker-free, motion-corrected cryo-electron tomographic alignment and reconstruction. Journal of Structural Biology: X 6, 100068 (2022).

55. Liu, Y.-T. et al. Isotropic reconstruction for electron tomography with deep learning. Nat Commun 13, 6482 (2022).

56. Wiedemann, S. & Heckel, R. A deep learning method for simultaneous denoising and missing wedge reconstruction in cryogenic electron tomography. Nat Commun 15, 8255 (2024).

57. Lamm, L. et al. MemBrain: A deep learning-aided pipeline for detection of membrane proteins in Cryo-electron tomograms. Computer Methods and Programs in Biomedicine 224, 106990 (2022).

58. Kremer, J. R., Mastronarde, D. N. & McIntosh, J. R. Computer Visualization of Three-Dimensional Image Data Using IMOD. Journal of Structural Biology 116, 71–76 (1996).

59. Jumper, J. et al. Highly accurate protein structure prediction with AlphaFold. Nature 596, 583–589 (2021).

60. Evans, R. et al. Protein complex prediction with AlphaFold-Multimer. 2021.10.04.463034 Preprint at 10.1101/2021.10.04.463034 (2022).

61. Pei, J., Tang, M. & Grishin, N. V. PROMALS3D web server for accurate multiple protein sequence and structure alignments. Nucleic Acids Res 36, W30–W34 (2008).

